# Competence repression by the SPβ prophage in *Bacillus subtilis* revealed by transcriptomics across culture conditions

**DOI:** 10.64898/2026.11.03.763063

**Authors:** Virginie Grosboillot, Luka Gnidovec, Špela Baebler, Jernej Ule, Anna Dragoš

## Abstract

Prophages are near-ubiquitous in bacterial genomes, yet their effects on host physiology are typically assayed under a narrow range of laboratory conditions, mostly involving DNA-damaging agents or physical and chemical stress. This leaves open the question of which ecological and physiological conditions trigger prophage-host interaction, and whether the outcome benefits the host or the prophage. Here, we used publicly available transcriptomic data to investigate conditions promoting prophage-host interaction in *Bacillus subtilis*. This analysis identified that competence strongly induces prophage gene expression. To further investigate the role of competence, we used two sets of *B. subtilis* strains: a natural SPβ lysogen strain and its cured counterpart, and a natural non-lysogenic strain and the same strain lysogenised with SPβ. Under competence-inducing conditions, both lysogen strains showed lower expression of the competence genes compared with their non-lysogenic counterparts. This result was corroborated by transformation assays that showed consistently lower transformation efficiency in the lysogen strains, and by fluorescence microscopy using a transcriptional reporter for the late competence gene. Because competence is a principal route for horizontal gene transfer in *B. subtilis*, prophage repression of the competence programme constrains host DNA uptake and evolvability. More broadly, these results show that mining public transcriptomes for condition effects can guide experimental effort towards prophage-host phenotypes that conventional laboratory conditions would miss.

## Introduction

Most sequenced bacterial genomes harbour at least one prophage, and in many species lysogeny is the rule rather than the exception: the prophage replicates as part of the host chromosome and is inherited vertically by the daughter cells (Dragoš et al., 2021b; Stefanic et al., 2025). These resident viral elements exert diverse and often unexpected effects on host physiology, ranging from the provision of virulence factors such as Shiga and cholera toxins (O’Brien et al., 1984; Waldor and Mekalanos, 1996), through bacteriocin production (Dragoš et al., 2021b), to population-level consequences of spontaneous induction (Toyofuku et al., 2017) and alterations of host cell biology, including cell shape and susceptibility to cell wall-targeting antimicrobials (Floccari et al., 2025). Resolving which effects a given prophage confers is crucial not only to fundamental microbial ecology, but also to applied settings: a beneficial strain may lose its desirable properties upon lysogenisation, or an otherwise commensal strain may acquire virulence traits. Importantly, the effects that prophages exert on their hosts are strongly condition-dependent, and both the environments a host encounters during its life cycle and the conditions under which a strain is deployed in industrial application vary widely.

Previous studies have examined how various stress-inducing conditions shift the balance between lysogeny and lysis (Sutcliffe et al., 2021; Dahlman et al., 2025). Much less is known about conditions that do not trigger lysis and in which lysogeny remains stable, yet that may modulate phage-host interactions. With a few exceptions, in which prophages carry recognisable functional modules such as anti-phage defence systems (Rousset et al., 2022) or bacteriocins (Dragoš et al., 2021b), the impact of a prophage on its host is difficult to predict from the phage genome alone. This is largely because such effects are frequently mediated by morons, whose function – unlike that of the structural genes of phages – remains poorly characterised (Brüssow et al., 2004; Tsao et al., 2018).

Several large-scale transcriptomic studies have mapped how diverse conditions reshape bacterial gene regulation (Nicolas et al., 2012; Rychel et al., 2020), and comparable condition-dependent regulation is expected for prophages residing within those hosts, and hence for phage-host interactions. Prophage regions, however, have rarely been the explicit focus of such analyses. Here, we ask whether phage-host interactions could be predicted from existing transcriptomic resources. We capitalise on a large condition-dependent transcriptomic dataset generated for the model bacterium *Bacillus subtilis*, obtained from a laboratory strain that carries the inducible prophage SPβ (Nicolas et al., 2012).

Robust analysis of expression variation across the prophage genome and different host culture conditions points to competence medium – the medium in which the host becomes naturally competent – as a strong candidate condition for previously unrecognised phage-host dynamics. To test this prediction, we used a set of natural lysogens and their prophage-free counterparts and generated our own transcriptomic data under three conditions: competence medium; mitomycin C, as a condition representing lytic induction; and stationary-phase LB, representing stable lysogeny, a state in which SPβ lysis-lysogeny decisions are governed by accumulation of the arbitrium signal (Erez et al., 2017).

Despite extensive, host-background-dependent effects, we find that the presence of the prophage consistently lowers expression of genetic competence genes in both host backgrounds tested. We confirm this effect using transformation assays with genomic and plasmid DNA, and a fluorescent P*_comGA_*-*gfp* reporter, a single-cell marker of late competence gene expression. Our work establishes a workflow that can be applied to other bacterial species for uncovering new mechanisms of host control by phages. It further shows that the phage maintains a similar expression pattern irrespective of host background, whereas different hosts respond differently to its presence – pointing to an asymmetry in regulatory control that favours the prophage.

## Materials and Methods

### Analysis of the transcriptomic dataset for condition selection and prophage genes of interest

For this analysis, we re-used the processed microarray dataset from Nicolas and coworkers (2012). The dataset was downloaded from the GEO repository (Accession number: GSE27219). The analysis was run with uv 0.12.19 and Python 3.13.15. The dataset was parsed with GEOparse 2.0.4 into a gene-annotation table (header.parquet; ID, ORF, GeneSymbol, GeneType) and an expression matrix (gene.parquet; 244 gene columns: 2847–3090, log_2_-scale intensities). Data quality was assessed with cuallee 0.15.4, including completeness of all variables and uniqueness of the ID, ORF and GeneSymbol identifiers. The analysis was limited to the prophage genes (ID: 2847 (*yotN*/*sprB*) to 3090 (*yokA*/*sprA*) inclusive).

#### Statistical analysis

All analyses were run in Python 3.13 (via uv 0.12.19) using NumPy 2.5.3, SciPy 1.18.1, statsmodels 0.15.0, scikit-learn 1.9.1, pandas 3.0.6 and DuckDB 1.5.6. Multiple-testing correction was performed using the Benjamini–Hochberg false-discovery rate (BH-FDR). Technical replicates were averaged within their biological replicate. Mean, median, standard deviation (SD) and variance ratio were calculated with NumPy and pandas (ddof = 1). To assess inter-condition variability, we identified within-condition replicate outliers from leave-one-out (LOO) residuals scaled by a robust per-gene σ (1.4826 × MAD), with two-sided normal p-values and BH correction (flagged at |z| > 4 and q < 0.05); each sample’s global offset (mean LOO residual) and replicate discordance D^2^ = mean (LOO residual^2^/σ^2^) were converted to modified z-scores (Iglewicz– Hoaglin). Condition responsiveness was tested by one-way ANOVA per gene with condition as the factor (between-group df = 101, within-group df = 156), reporting the F-statistic, the effect size η^2^, and BH-adjusted q-values. Because variance scales with mean expression, we fitted a LOWESS mean–variance trend and tested for excess total variance using the robust z (median/MAD) of the trend residual (one-sided normal test, BH). The signal-to-noise ratio was defined as the ratio of between-condition SD to within-condition SD.

#### Visualisation

Results were plotted using matplotlib 3.11.2, seaborn 0.13.2 and adjustText 1.4.0. For the heatmap and gene ranking, each condition was summarised as the mean of its biological replicates. Only conditions with biological replicates ≥ 2 were used for the heatmap.

### Bacterial strains, media and culture conditions

The bacterial strains and plasmids used in this study are listed in Table 1.

**Table 1.** Bacterial strains and plasmids used in this study. Cm^R^: chloramphenicol resistance, Amp^R^: ampicillin resistance and Spec^R^: spectinomycin resistance

| Strain | Relevant genotype / description | Source |
|---|---|---|
| ***B. subtilis*** |  |  |
| 168 | WT / lysogen for SPβ | Kunst et al. (1997) |
| 168 | P*comGA-gfp*, Cm<sup>R</sup> | Smits et al. (2005) |
| 3610 (DS215) | *swrA*::*tet* | Kearns et al. (2004) |
| Δ6 | ΔSPβ Δskin ΔPBSX Δprophage1 *pks*::*cat* Δprophage3, Cm<sup>R</sup> | Westers (2003) |
| MB8_B7 | *amyE*::P<sub>pL</sub>-*gfp*, Cm<sup>R</sup> | *This work* |
| MB8_B7 | WT / lysogen for SPβ | Kiesewalter et al. (2021) |
| MB8_B7 ΔSPβ | SPβ::*erm* | Dragoš et al. (2021a) |
| P9_B1 | WT / non-lysogen | Kiesewalter et al. (2021) |
| P9_B1 (DTUB43) | *amyE*::P<sub>hyper-spank</sub>-*sfGFP*, Cm<sup>R</sup> | Dragoš et al. (2021a) |
| P9_B1 lysSPβ (DTUB231) | lysogen for SPβ | Dragoš et al. (2021a) |
| P9_B1 lysSPβ P<sub>*comGA*</sub>-*gfp* | P<sub>*comGA*</sub>-*gfp*, Cm<sup>R</sup> | *This work* |
| P9_B1 P<sub>*comGA*</sub>-*gfp* | P<sub>*comGA*</sub>-*gfp*, Cm<sup>R</sup> | *This work* |
| ***E. coli*** |  |  |
| DH5α | F<sup>-</sup> *φ80dlacZΔM*15, Δ(argF-lac)169, U169, deoR, *rec*A1, *end*A1, hsdR17(rK<sup>-</sup>mK<sup>+</sup>), phoA, supE44, λ<sup>-</sup>, thi-1, *gyr*A96, relA1 | Woodcock et al. (1989) |
| DH5α | pTB233-P<sub>*comGA*</sub>-*gfp* | *This work* |
| MC1061 | P<sub>hyper-spank</sub>-*sfGFP*, Cm<sup>R</sup> | Van Gestel et al. (2014) |
| **Plasmids** |  |  |
| pTB233 | pTB233, Amp<sup>R</sup>/Spec<sup>R</sup> | Gallegos-Monterrosa et al. (2021) |
| pGFP-rrnB | *amyE*, P<sub>rrnB</sub>-*gfp*, Spec<sup>R</sup> | Veening et al. (2009) |

For the RNA-seq study, *Bacillus subtilis* strains were grown overnight in Luria–Bertani (LB) broth at 37°C with shaking at 220 rpm. Growth conditions were adapted from Nicolas et al. (2012). Briefly, for the LBstat and M40t90 conditions, cells were grown overnight in LB broth at 37°C with shaking at 220 rpm. Overnight cultures were diluted 2000-fold in fresh pre-warmed LB broth. Cells were grown at 37°C with shaking at 220 rpm and cell biomass was monitored with a cell growth quantifier (CGQ, Scientific Bioprocessing). For LBstat, cells were collected during stationary phase. For M40t90, 40 ng/mL mitomycin C was added during exponential phase; cells were collected after 90 min. For C90, cells were plated on LB agar and incubated overnight at 30°C. The colonies were used to inoculate pre-warmed SPI medium to an OD_600_ of 0.5 and cultures were grown at 37°C with shaking at 220 rpm. Cell biomass was monitored with the CGQ and when the growth curve departed from the exponential phase, the cells were diluted tenfold (1:10) in fresh pre-warmed SPII medium and incubated for an additional 90 min. Three biological replicates per condition were performed, and the samples are listed in Table S1.

For gDNA and plasmid DNA extraction, the respective strains were grown overnight in LB broth supplemented with the appropriate antibiotics: 25 µg/mL chloramphenicol for *E. coli*, and 5 µg/mL chloramphenicol and 10 µg/mL tetracycline for *B. subtilis*. Plasmid DNA was extracted with the QIAprep Spin Miniprep Kit (QIAGEN) and gDNA with the Bacterial and Yeast Genomic DNA Purification Kit (EURx). Concentrations were measured with NanoDrop and samples were stored at -20 °C until use.

For the transformation assay, *B. subtilis* was grown in Modified Competence (MC) medium (Chastanet, 2009, *unpublished laboratory protocol*), based on the classical medium of Anagnostopoulos and Spizizen (1961). The MC medium (10×) was prepared with 14 g K_2_HPO_4_·3H_2_O, 5.2 g KH_2_PO_4_, 20 g glucose, 10 mL of trisodium citrate (300 mM), 1 mL ferric ammonium citrate (2.2 %), 1 g casein hydrolysate, 2 g potassium glutamate and ddH_2_O to a final volume of 100 mL. Competent cultures were grown in 2 mL 1× MC, supplemented with 6.7 µL MgSO_4_ (1 M).

For the plaque assay, *B. subtilis* Δ6 was grown overnight in LB broth at 37°C with shaking at 220 rpm.

### Host cell and phage particle concentrations

For each sample, bacterial and phage concentrations were measured at sampling time. Colony-forming unit (CFU) was determined as follows: serial 1:10 dilutions of the cultures were prepared in 0.9 % NaCl and 100 µL of the dilution was plated on LB plates. Dilutions were plated in triplicate and the concentration was determined as the mean bacterial CFU/mL. Plaque-forming unit (PFU) was determined with the double agar overlay assay (Kropinski et al., 2009). A 2 mL culture volume was centrifuged at 8,000 × *g* for 3 min. Supernatant was filtered with a 0.2 µm filter and 100 µl of serial 1:10 dilutions of the filtrate was used and mixed 1:1 (v/v) with Δ6 culture and 3 mL pre-warmed LB 0.3 % soft agar. Three dilutions were plated for each sample.

### RNA extraction

RNA extraction was performed according to protocols 4 and 7 of the RNeasy Mini Kit (QIAGEN) with minor modifications. After addition of the sample at a 1:3 (v/v) ratio to the RNAprotect Bacteria Reagent (QIAGEN), the cells were washed twice with 1 mL of RNAprotect, resuspended and centrifuged for 1 min at 5000 × *g*. After removal of the fixative, the cells were stored at -80 °C until processing. For the extraction, 18 µL of lysozyme (250 mg/mL in nuclease-free water) and 100 µL of proteinase K (20 mg/mL, Thermo Fisher) were added to each sample. Samples were mixed thoroughly by pipetting and incubated at room temperature for 10 min. During incubation, samples were vortexed for 10 s every 2 min. The volumes of RLT buffer and absolute ethanol were adjusted to 413 µL and 295 µL, respectively. The wash with 700 µL of RW1 buffer was performed twice and the wash with 500 µL of RPE buffer was performed three times, transferring the column to a new 2 mL tube each time. Centrifugations were carried out for 30–60 s at maximum speed for all washing steps. The final centrifugation step was performed for 5 min at maximum speed to remove residual ethanol. Columns were then transferred to a new 1.5 mL tube; 45 µL of pre-heated (80°C) nuclease-free water was added to the column membranes and incubated for 10 min at room temperature. Columns were centrifuged for 2 min at maximum speed. To remove genomic DNA present in the samples, DNase treatment was performed with TURBO DNase (TURBO DNA-free™ Kit, Invitrogen) in a total volume of 100 µL using 2 µL of DNase and the two-step incubation protocol. To remove the DNase, a second round of RNA clean-up was performed on the RNeasy Kit columns: 250 µL of absolute ethanol was mixed with 100 µL of sample and the total volume was loaded onto the column. Two washes were performed with the RPE buffer and RNA was eluted in 35 µL nuclease-free water. RNA concentrations were measured with NanoDrop and samples with low 260/230 ratios were subjected to a new round of RNA clean-up. Samples were then run on a 1 % (w/v) TAE agarose gel to check the integrity of the extracted RNA.

### Library preparation

Sequencing libraries were prepared using a template-switching oligonucleotide (TSO)-based protocol adapted from Faraway et al. (2025). RNA fragmentation was performed by adding 0.5 µL of 5× SuperScript IV (SSIV, Invitrogen) buffer to 3.5 µL of RNA (105 ng), followed by incubation at 95°C for 4 min. Fragmented RNA was mixed with 10.5 µL of reverse transcription (RT) buffer (60 mM KCl, 10 mM MgCl_2_, 3.5 mM dNTPs, 6 mM DTT, 2 µM randomer_RT primer containing Illumina P7 sequences, 2.5 µM TSO primer containing Illumina P5 sequences and unique molecular identifiers (UMIs), 10 mM Tris-HCl pH 8.0) and incubated for 3 min at 65°C and then cooled to 42°C at a rate of 0.5°C/s. A mixture of 0.25 µL RNasin Plus RNase Inhibitor and 0.25 µL SSIV was added to the samples while keeping them at 42°C, followed by incubation at 42°C for 90 min. The RT product was purified using 1.2× volume of NGS TotalPure beads and resuspended in 10 µL nuclease-free water. Indexing PCR was performed in 40 µL (Q5 2× PCR master mix, 0.25 µM P7/P5 Illumina indexed primer mix) using 4 µL of eluted cDNA (PCR program: 98°C for 40 s; 7 cycles of 98°C for 20 s, 65°C for 30 s and 72°C for 45 s; 72°C for 3 min). A 6 % TBE gel (Thermo Fisher) was run with 5 µL of each PCR product (30 min, 180 V) and stained with SYBR Green I for 30 min to verify successful amplification of each sample before pooling. The PCR products were pooled based on the library band intensities on the gel to achieve approximately equal representation of each library. The pooled libraries were purified using 0.8× volume of NGS TotalPure beads and eluted in 20 µL water. The final concentration was measured with Qubit. All samples were processed at the same time. The correspondance between P7/P5 Illumina indexed primer and samples is listed in Table S1 and all primer sequences are listed in Table S2.

### rRNA depletion

The depletion of rRNA was performed according to Wilkins and Ule (2021). The list of generated sgRNAs for *B. subtilis* rRNA depletion is in Table S3. The re-amplification of the library was performed with 9 cycles and the P5_multiplex_primer/P7_multiplex_primer pair (Table S2). The PCR was cleaned up using 0.8× volume of NGS TotalPure beads and eluted in 20 µL water. Final library concentration was measured with Qubit.

### RNA-seq data analysis

The complete genome sequences for *B. subtilis* 168 (NCBI, Accession Version: NC_000964.3; Borriss et al. (2018)), *B. subtilis* MB8_B7 (NCBI, Accession Version: CP045821.1; Kiesewalter et al. (2021)) and *B. subtilis* P9_B1 lysSPβ (NCBI Sequence Read Archive, Accession numbers SRX26270955 and SRX26270956; Dragoš et al. (2021a)) were re-annotated and used for the analysis. Each of the three *Bacillus* genomes was split at the SPβ attachment site (attP/attB: ACAGATAAAGCTGTAT, Kohm et al. (2022)) into a bacterial contig and a prophage contig. The bacterial contig was re-annotated with Bakta 1.12.1 (database v6.0, full). The prophage contig was annotated with Pharokka 1.10.1 (prodigal-gv, --dnaapler) and functionally refined with phold 1.2.6. The two contigs form a combined 2-contig reference per strain (4,212 / 4,185 / 4,184 CDS for 168 / MB8_B7 / P9_B1 lysSPβ, respectively) used for read mapping and quantification. Samples were mapped to the reference of their own strain. For between-strain prophage comparisons, only the 167 consensus genes (one-to-one orthologues present in all three prophages) were retained (Table S4 and Figure S1).

Sequencing reads were processed as follows. UMIs were extracted with umi-tools 1.1.5 (--bc-pattern=NNNNCCCCNNN). Reads were trimmed with fastp 0.23.4 (5’ front trimming of 3 bases; minimum read length 20 bp). Reads were mapped with bowtie2 2.5.4 (end-to-end) and alignments were sorted and indexed with samtools 1.22.1. PCR duplicates were then collapsed with umi_tools dedup --paired (default error-aware directional UMI clustering; UMIs (7-nt random segments) were on the TSO primer (read1)) and the deduplicated BAMs were indexed with pysam 0.24.1 (htslib BAI); counts therefore represent unique molecules rather than PCR copies. Gene-level counts were obtained with featureCounts (subread 2.1.1; -p -O -t CDS -g transcript_id) as integer counts on CDS features. The -O flag assigns a multi-overlap fragment to every overlapped feature (liberal counting for operons). Bacterial and prophage genes were separated according to the contig/locus-tag scheme of each combined reference.

Differential expression was performed with PyDESeq2 0.5.4 (negative binomial generalised linear model, Wald test, Benjamini–Hochberg false-discovery rate). The design was a single-factor model (∼has_prophage, ∼condition or ∼strain, depending on the comparison) with the reference level set explicitly. Unless stated otherwise, genes with padj < 0.05 were considered differentially expressed and log_2_ fold-changes (log_2_FC) were reported with apeglm shrinkage. To define a high-confidence core signature, a second filter of |log_2_FC| > 2 (shrunken) was applied for the prophages, because they carry few genes. Independent quantification with FADU (fractional, operon-aware) was used as a sensitivity check and agreed with featureCounts on the UMI-deduplicated counts (Pearson *r* = 0.70–0.999, median 0.91; Spearman *r* = 0.87–0.996, median 0.96 per sample; the spread reflects the ≈2.5-fold lower molecule count after deduplication). Principal-component analysis of variance-stabilised counts was used to assess sample grouping. Gene Ontology (GO) and COG over-representation tests (hypergeometric, Benjamini–Hochberg) were run on Bakta annotations for bacterial gene sets. GO identifiers and names were resolved against the Gene Ontology release 2026-07-26 (go.obo, full ontology; go-basic.obo, same release, for term names) to update obsolete identifiers.

### Transformation assay

For the transformation assay, we used the simplified one-step protocol in MC medium according to a modified Losick laboratory protocol (Chastanet (2009), unpublished laboratory protocol). Briefly, 20 µL of the overnight culture was transferred to 2 mL of MC medium and incubated for 6 h at 37°C with shaking at 220 rpm, to allow competence to develop. The culture was then split into two 1 mL cultures: 1 µg of DNA was added to one of the cultures and 1 µL nuclease-free water was added to the other culture as a control. Cultures were incubated at 37°C with shaking at 220 rpm. After 30 min, 1 mL of pre-warmed LB medium was added to the cultures and cells were incubated for an additional 90 min. Subsequently, serial 1:10 dilutions of the cultures were prepared in 0.9 % NaCl and 100 µL of the dilution was plated on LB plates without antibiotics using glass beads. To enumerate transformants, cultures were plated undiluted on selective LB plates containing either 10 µg/mL tetracycline (transformation with *swrA*::*tet* gDNA) or 100 µg/mL spectinomycin (transformation with gDNAs: *amyE*::P_pL_-*gfp* and *amyE*::P_hyper-spank_-*sfGFP*, and plasmid: P_hyper-spank_-*sfGFP*). All dilutions and cultures were plated in triplicate to account for plating errors.

All transformation assays were performed with three biological replicates per strain. Transformation frequency was determined as the ratio of transformant colony-forming units (TCFU) to total viable cells and was min–max normalised per DNA type and day to enable visual comparison of the lysogens versus non-lysogens for the different conditions tested on the same plot. Statistical significance between lysogens and non-lysogens was determined using a two-sided Welch t-test on log_10_-transformed transformation frequency, as transformation frequency is approximately log-normally distributed, for each experiment.

### Strain and fluorescence reporter construction

For the transformation assay, *B. subtilis* MB8_B7 *amyE*::P_pL_-*gfp* strain was obtained as follows. The P_pL_ promoter region was amplified from genomic DNA of *B. subtilis* 168 by PCR using primers oAD10/oAD11 (Table S5), which introduced EcoRI and NheI restriction sites, respectively. The amplified promoter fragment was cloned into the pGFP-rrnB plasmid, which carries homologous regions for integration at the *amyE* locus and a chloramphenicol resistance marker. Both the purified PCR product and the pGFP-rrnB plasmid were digested with EcoRI and NheI, followed by ligation. The resulting P_pL_-*gfp* plasmid was verified by gel electrophoresis and Sanger sequencing. *B. subtilis* MB8_B7 was transformed with the P_pL_-*gfp* plasmid using the modified Losick protocol (see Transformation assay section) and transformants were selected based on chloramphenicol resistance. Integration of the construct at the *amyE* locus was confirmed by PCR using the primers amyEF/amyER. Materials and Methods To create the late competence gene reporter strains, the upstream DNA fragment containing the *comGA* promoter and the *gfp* gene and the downstream DNA fragment containing the *comGA* promoter and the *comGA* gene were PCR-amplified from genomic DNA of *B. subtilis* 168 P*_comGA_*-*gfp* with oVG126/oVG128 and oVG130/oVG127, respectively (Table S5). The digested DNA fragments (SacI/BamHI and HindIII/SalI) were sequentially cloned into plasmid pTB233 containing a spectinomycin resistance cassette. The resulting plasmid pTB233-P*_comGA_*-*gfp* contains the spectinomycin resistance cassette flanked by the upstream and downstream regions. The plasmid was cloned and maintained in *E. coli* DH5α. The correct construct was confirmed by sequencing. *B. subtilis* reporter strains were obtained via natural competence transformation with pTB233-P*_comGA_*-*gfp*, following the protocol described in the Transformation assay section. LB plates supplemented with 100 µg/mL spectinomycin were used for selection.

### Fluorescence microscopy

Cells of the wild-type and SPβ strains carrying the P*_comGA_*-*gfp* transcriptional reporter were grown as described in the Transformation assay section. At 6 h and 8 h of incubation, samples were collected and mounted on 1.5 % agarose pads in LB. Images were acquired on a Zeiss Elyra 7 microscope controlled by ZEN black software (v16.0), using a Plan-Apochromat 63×/1.4 Oil DIC M27 objective and a pco.edge 4.2 sCMOS camera (16-bit, 1×1 binning, gain 1), giving a pixel size of 0.0968 µm. For each field, a transmitted-light brightfield image and a GFP fluorescence image were recorded sequentially. GFP was excited with a 488 nm diode laser (HR Diode 488–100; 0.1 % laser power) in wide-field epifluorescence mode through a quad-band LBF 405/488/561/642 filter, with 50 ms exposure. Acquisition settings were identical for all samples. Larger areas were imaged as tile scans of up to 4 × 4 fields (1280 × 1280 px each, ≈3 % overlap), stitched in ZEN, in addition to single fields. Three biological replicates were imaged per strain and time point, with one or two tile scans per replicate.

### Image analysis

Images were analysed using custom Python 3.12 scripts with pylibCZIrw 6.1.0, Cellpose 3.1.1 (Stringer et al., 2021; Stringer and Pachitariu, 2025), scikit-image 0.26.0, SciPy 1.18.1 and NumPy 2.0.2. Brightfield and GFP images were aligned using a constant channel offset determined from 514 isolated GFP-positive cells. To ensure reliable cell detection, tilescan regions were screened for image quality based on cell focus, edge sharpness and cell number. Only regions meeting predefined quality criteria (≥ 80 % in-focus cells, edge steepness ≥ 0.30 and ≥ 10 cells) were included, and overlapping regions were excluded. Cells were segmented from brightfield images independently of GFP fluorescence using Cellpose (bact_phase_cp3, diameter 20 px). To account for differences in cell contrast, segmentation was performed on both original and contrast-inverted images, and the results were combined. An additional image-processing step was used to detect isolated rod-shaped cells missed by Cellpose. Segmented objects were filtered by size, shape and cell contacts to exclude aggregates, irregular objects and cells touching the image borders. Valid cells had an area of 0.6–10 µm², a width of 0.45–2.0 µm and a solidity ≥ 0.60. Cells in short chains were retained, whereas larger clusters were excluded. GFP fluorescence was quantified for each cell after local background subtraction. Relative fluorescence was calculated as ΔF/F = (mean cellular GFP intensity − local background)/local background. Fluorescence immediately surrounding each cell was also measured to distinguish cell-associated GFP signals from background fluorescence or signals originating from neighbouring cells. Cells were classified as GFP-positive when ΔF/F > 0.30 and the difference between cellular and surrounding fluorescence was ≥ 0.10. These thresholds were validated by visual inspection. The percentage of GFP-positive cells was calculated relative to the total number of valid cells for each biological replicate and condition. In total, 3,697 (WT, 6 h), 2,422 (SPβ, 6 h), 3,100 (WT, 8 h) and 600 (SPβ, 8 h) cells were analysed. Representative images show 62 × 62 µm regions selected from well-focused areas. Identical display settings were used across conditions for each imaging channel (brightfield: 470–5700 grey values; GFP: a fixed linear intensity range). Images were processed in Fiji/ImageJ 2.18.0/1.54p (Schindelin et al., 2012) to adjust brightfield brightness and add scale bars to the images.

## Results

### Most SPβ prophage genes vary little across culture conditions, but a small subset is strongly condition-responsive

We used the published transcriptomic dataset from Nicolas and coworkers (2012) to identify conditions of interest for studying the impact of SPβ prophage on *B. subtilis* physiology. The dataset includes SPβ prophage, which is integrated in the *B. subtilis* 168 genome, and covers 102 culture conditions (full list in Table S6). It is therefore a rich source of information for selecting conditions of interest, even though natural strains may behave differently from the laboratory strain *B. subtilis* 168.

After initial data structure assessment (Supplementary Results S1), we then assessed the reproducibility between each replicate for a given culture condition. Only two conditions, S8 (n = 2) and M40t90 (n = 3), showed high within-condition variability among biological replicates (Figure 1 A). As S8 had only two biological replicates, we could not determine which one caused the dispersion, while for the mitomycin C condition, it was associated with replicate M40t90_2 (Figure 1 B). To identify the conditions affecting SPβ prophage gene expression, we then examined its expression profile across all samples. Most pro-phage genes have relatively low expression (Figure 1 C), with half of the samples having a log_2_ intensity below 8.7. We also observed that most genes had a consistent expression level across conditions, with a median log_2_ intensity SD of 0.84 and only 77 genes above 1.0 (Figure 1 D). Four genes showed particularly high variability across conditions, with SD above 2.0 (*yoqM*: 2.27, *sspC*: 2.17, *bdbA*: 2.05, *sunT*: 2.04) and 13 between 1.5 and 2.0 including *yolJ* (1.87), *yotN* (1.76), *bdbB* (1.76), *yolA* (1.63), *yolB* (1.60) and *yosA* (1.59). To verify that the dispersion is real and not due to intra-condition variability or high expression levels, we plotted the SD across all replicates and conditions against gene-wise mean expression, fitted a LOWESS trend, and tested condition responsiveness (Figure 1 E). Only *yoqM*, *sspC*, *yotN* and *yosA* displayed a significant variance in log_2_ expression intensity across conditions (BH q < 0.05). The fitted LOWESS trend also showed that SD increases with gene mean expression, so a high SD alone is not evidence of hypervariability. Accordingly, the variance of *bdbA* and *sunT* was not significant despite a high calculated SD (> 2.0). The signal-to-noise analysis confirmed this classification. The *yolJ* gene was additionally classified as highly variable with low noise; *bdbA*, *sunT* and surrounding genes in the genome were highlighted as highly noisy and therefore not biologically relevant (Figure 1 F).

**Figure 1.**
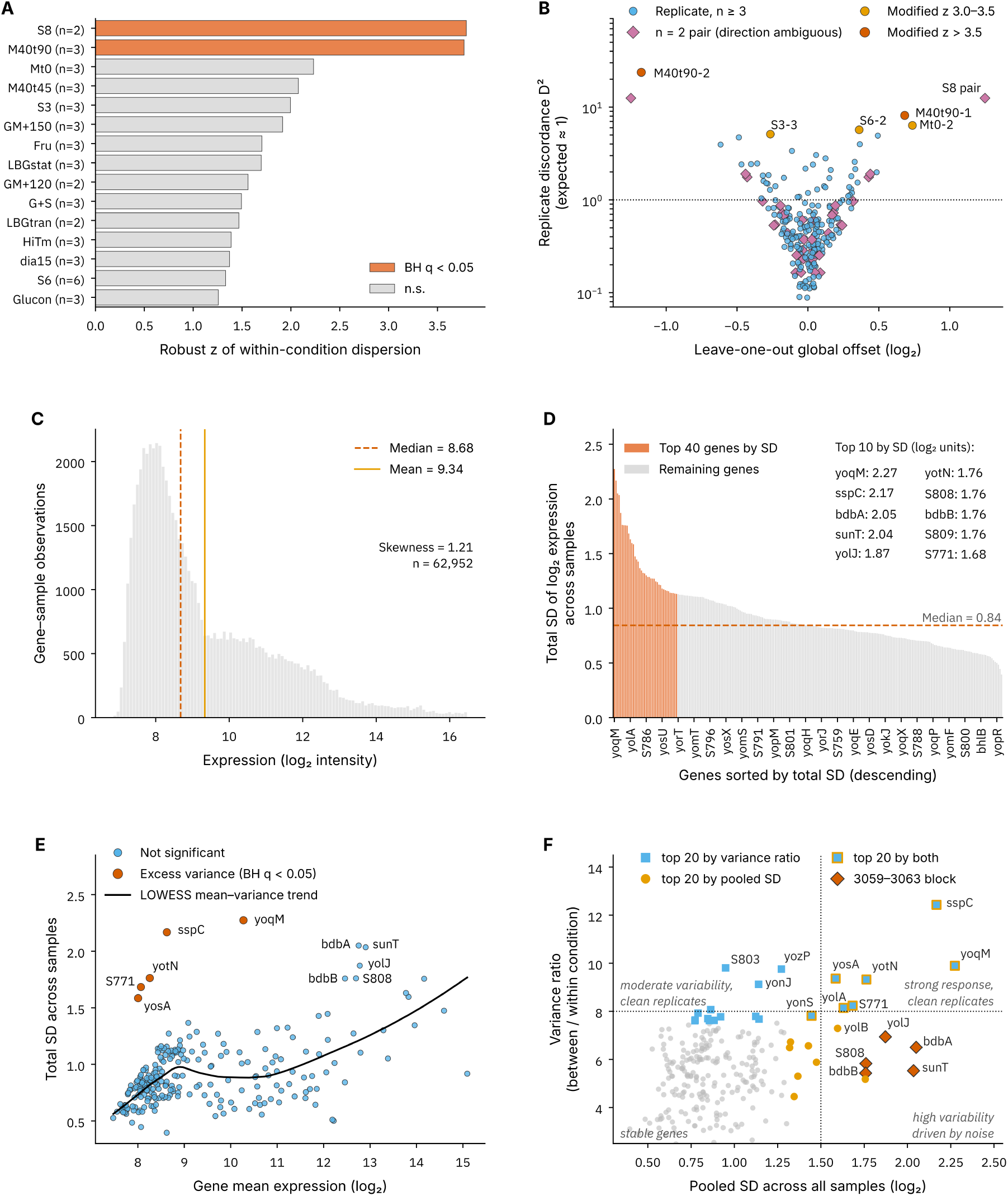
Gene expression (log_2_ intensity) variation across conditions and SPβ prophage genes. (A) Top-15 conditions ranked by the robust z of their within-condition dispersion. For each condition (n ≥ 2 biological replicates), replicate scatter per gene was normalised by the pooled within-gene variance and summarised as a dispersion ratio; the robust z (modified z, median/MAD) of the log ratio is plotted. Bars are coloured by Benjamini–Hochberg-adjusted significance (q < 0.05, orange; not significant (n.s.), light grey). n = number of biological replicates. S8 and M40t90 are the only significantly over-dispersed conditions. (B) Replicate discordance versus leave-one-out global offset. Each point represents a biological replicate (for conditions with n ≥ 2). The global offset represents the mean log_2_ shift of the sample relative to the mean of the other replicates in its condition; the replicate discordance D^2^ is the mean squared leave-one-out residual across genes divided by the pooled within-gene variance (expected ≈ 1, dotted line). Colours encode the modified z of D^2^ (> 3.5, orange; 3.0–3.5, yellow; otherwise blue); n = 2 pairs are shown as pink diamonds to mark that the direction of discordance is ambiguous for those replicates. (C) Distribution of log_2_ intensity across samples for each prophage gene. The mean and the median are indicated with yellow and orange lines respectively. (D) Gene expression standard deviation across conditions, ordered by descending SD values. The median SD value is indicated with a dotted orange line. In orange are the top-40 genes with highest SD values (an arbitrary cut-off). For clarity, the names of the 244 genes were omitted but the top-10 genes ranked by SD were listed in the figure. (E) Total standard deviation (SD) across all biological-replicate samples versus gene mean expression for all genes. The black curve is the LOWESS mean-variance fitted curve (frac = 0.5). Genes whose variance significantly exceeds the trend expectation are shown in orange and labelled (robust z of the trend residual, one-sided normal test, Benjamini–Hochberg q < 0.05); all other genes are shown in blue. The genes showing a high SD for which the variance is not significant are also labelled. (F) Signal-to-noise landscape of gene expression. Each point represents a gene. The variance ratio V = σ_between_ / σ_within_, where σ_between_ is the standard deviation of the condition means and σ_within_ is the median within-condition standard deviation (conditions with n ≥ 2). High V indicates a gene whose expression changes strongly across conditions while its replicates stay concordant. Blue squares mark the top-20 genes by variance ratio, yellow circles the top 20 by pooled SD, and blue points with an orange edge are in both top-20 sets. Orange diamonds mark the 3059–3063 block (S808, *bdbB*, *yolJ*, *bdbA*, *sunT*), which has high pooled SD but low signal-to-noise. Reference lines are drawn at variance ratio = 8 and pooled SD = 1.5.

Overall, this analysis identified a subset of prophage genes exhibiting greater expression variability across culture conditions than expected from their mean expression levels. This indicates that the expression of a limited number of prophage modules may be particularly sensitive to changes in culture conditions.

### Culture conditions fall into distinct prophage-response clusters and reveal candidates for lysogeny studies

Our next goal was to identify culture conditions with the strongest effects on the prophage gene expression profile. For this, we used the Δ log_2_, the log_2_ ratio of a condition’s mean expression to the mean across all conditions. Distinct patterns of higher (Δ log_2_ > 0) and lower (Δ log_2_ < 0) expression relative to the all-sample mean were observed across the conditions (Figure 2 A). Mitomycin C treatment (M40t90) was the only condition with a completely distinct pattern across the top-40 SD-ranked genes; two gene clusters lay well above the all-sample mean, whereas only a few genes lay below it. Swarming (Sw) and growth at high colony density on plates (BC) shared a distinct effect on the prophage: 36 genes of the 40 lay below their average expression across all conditions (Figure 2 A). We also noticed that sporulation (S) and biofilm (B) conditions shared a similar gene expression pattern, with only six genes well above the mean and more than 75 % of the genes below it. Stress conditions, such as growth at cold temperature (Cold), growth in competence medium (C90) or growth in the presence of ethanol (Etha) produced a weaker shift in the same direction as mitomycin C, with the exception of the first gene cluster (<10 genes) (Figure 2 A).

**Figure 2.**
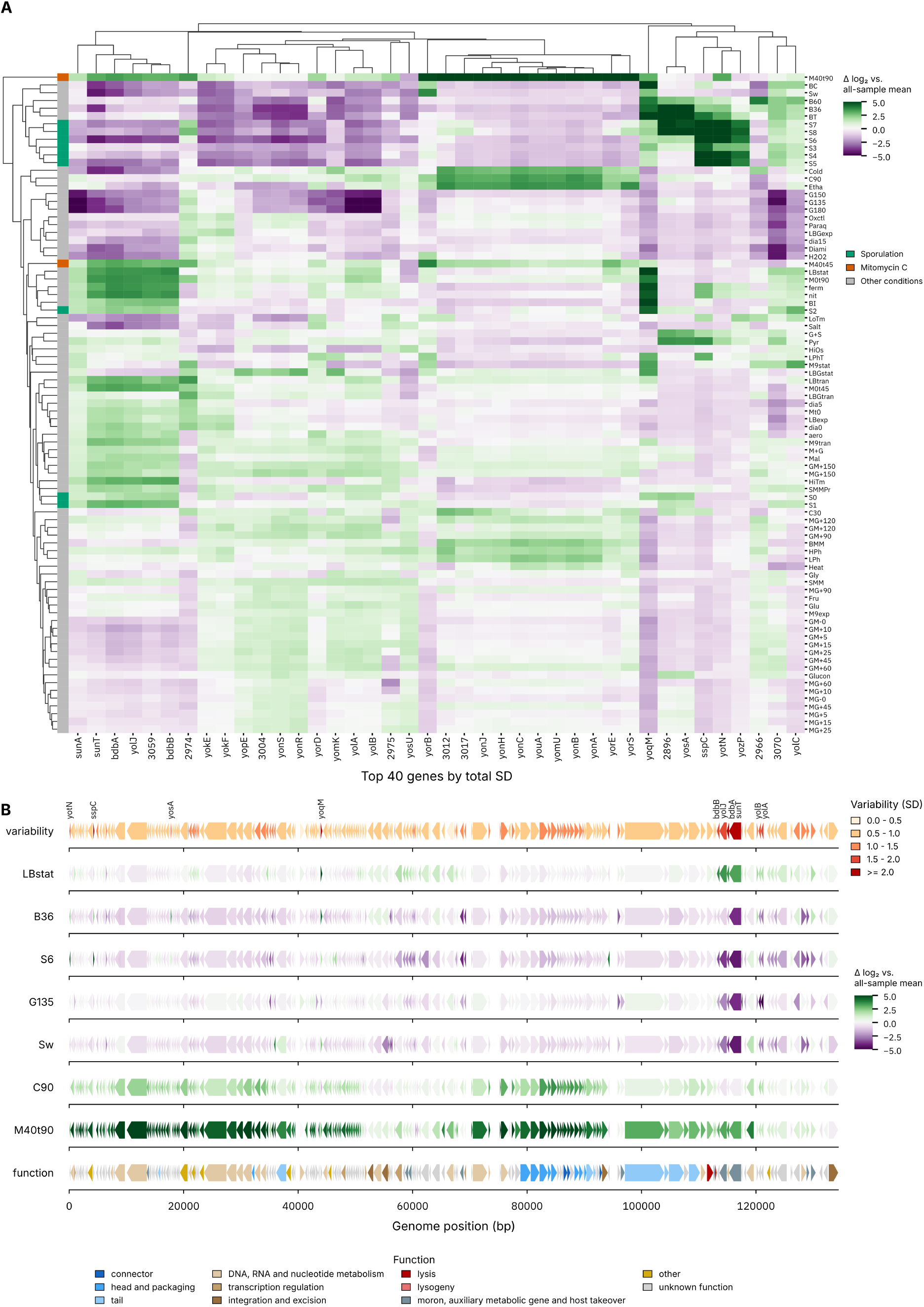
Prophage gene expression pattern across conditions. (A) Clustered heatmap of the 40 most variable genes across conditions. Each cell shows a condition’s mean expression relative to the all-sample mean (Δ log_2_; PRGn diverging colour scale, centred at 0, clipped at ±5). Genes (columns) are the 40 genes with the highest total standard deviation across all 258 biological-replicate samples; conditions with fewer than two biological replicates were excluded (85 conditions). Rows and columns are ordered by hierarchical clustering (Euclidean distance, average linkage). Branch heights have been re-scaled for display (monotonic power transform d to dp). The row-colour bar indicates the condition group: sporulation (S0–S8, green), mitomycin C (M40, orange), other conditions (grey). (B) Gene expression pattern across the full SPβ genome for seven selected conditions. The first track uses the SD calculated across all conditions to locate the genes with the highest variability. Genes with SD > 1.5 are annotated on the graph. The middle tracks represent the condition’s mean expression relative to the all-sample mean (Δ log_2_; PRGn diverging colour scale, centred at 0, clipped at ±5). The bottom track represents the gene functions – derived from the *B. subtilis* 168 prophage annotation file.

Based on the condition clusters, the transcriptomic response pattern and the intensity of the gene expression shifts, we reasoned that selecting one condition per well-defined cluster would allow exploration of the prophage-host relationship in very different environments, and might help to elucidate as yet unknown consequences of lysogeny and suggest functions for some uncharacterised prophage genes. Suggested conditions for further studies include: mitomycin C treatment for 90 min, swarming, biofilm after 36 h and sporulation after 6 h, growth in competence medium for 90 min, germination after 135 min and growth in LB to stationary phase (see Table 2).

**Table 2.** Conditions from Nicolas et al. (2012) considered of interest in this study.

| Condition ID | Description |
|---|---|
| B36 | biofilm formation_early biofilm = 36h of growth in MSgg medium without shaking |
| C90 | competence_time=90 minutes after induction |
| G135 | germination_time=135 minutes after induction |
| LBstat | growth in LB medium_stationary phase |
| M40t90 | growth in LB until exponential phase_90 minutes after addition of 40 ng/mL mitomycin |
| S6 | induction of sporulation_time=6h |
| Sw | 18h of growth on soft plates |

Finally, we extended this representation (condition mean expression relative to the all-sample mean) to all prophage genes for the selected conditions (Figure 2 B). The similarity pattern between C90 and M40t90 was maintained, with the Δ log_2_ value of most genes involved in replication and transcription well above the all-sample mean, as were genes encoding structural proteins (Figure 2 B). In the LBstat condition, Δ log_2_ was close to 0 for most genes, with a few genes lying above the all-sample mean, but unlike in C90 and M40t90, these genes did not encode phage structural proteins. For the remaining conditions, prophage gene expression was mostly below the all-sample mean (Figure 2 B), including genes involved in replication and structural protein production. For B36 and S6, we also observed that the excisionase *yotN* and several other uncharacterised genes lay well above the all-sample mean.

According to this analysis, SPβ gene expression is affected by its host growth conditions, and the effect is highly variable in terms of number of genes affected, direction of transcriptional shift and amplitude of the shift (Figure 2 A).

### Transcriptomic profiling of wild *B. subtilis* strains confirms that the prophage gene expression profile is affected by the host growth condition

Next, we extended the findings from the transcriptomic dataset of the laboratory strain *B. subtilis* 168 to search for new examples of phage-host interaction within natural *B. subtilis* isolates. We selected culture conditions that affected the pro-phage transcriptomic profile and were therefore relevant to phage-host interactions (Figure 2). We tested their effects in two natural *B. subtilis* strains representing distinct, but related genetic backgrounds (Table S7). Including two different host backgrounds allowed us to assess the broader applicability and robustness of our initial predictions.

We therefore generated our own RNA-seq data for the conditions selected above. We chose growth in competence medium for 90 min (C90) and growth in LB with 40 ng/mL mitomycin C for 90 min (M40t90) because of their strong influence on the prophage transcriptomic pattern, and used growth in LB to stationary phase (LBstat) as a control condition. We applied these three conditions to four strains in two pairs – MB8_B7 and its ΔSPβ derivative, and P9_B1 and the same strain lysogenised with SPβ – and we also included *B. subtilis* 168, to allow direct comparison with the public dataset analysed above. Each condition was performed in three biological replicates (45 libraries in total; Table S1). Libraries were sequenced, mapped to strain-specific bacterial and prophage references, quantified at the CDS level and compared with PyDESeq2 as described in Methods; unless stated otherwise, genes with padj < 0.05 were considered differentially expressed, and a |log_2_FC| > 2 threshold was additionally applied to prophage genes.

We first compared prophage transcripts in lysogens grown in C90 or M40t90 versus LBstat (padj < 0.05) and we confirmed that the host growth condition affects prophage gene expression. There was a clear shift in gene expression between the conditions for the SPβ prophage in both MB8_B7 and P9_B1 strains (Figure 3 B, C, E and F), and the shift was more pronounced for these strains than for 168 (Figure 3 A and D).

**Figure 3.**
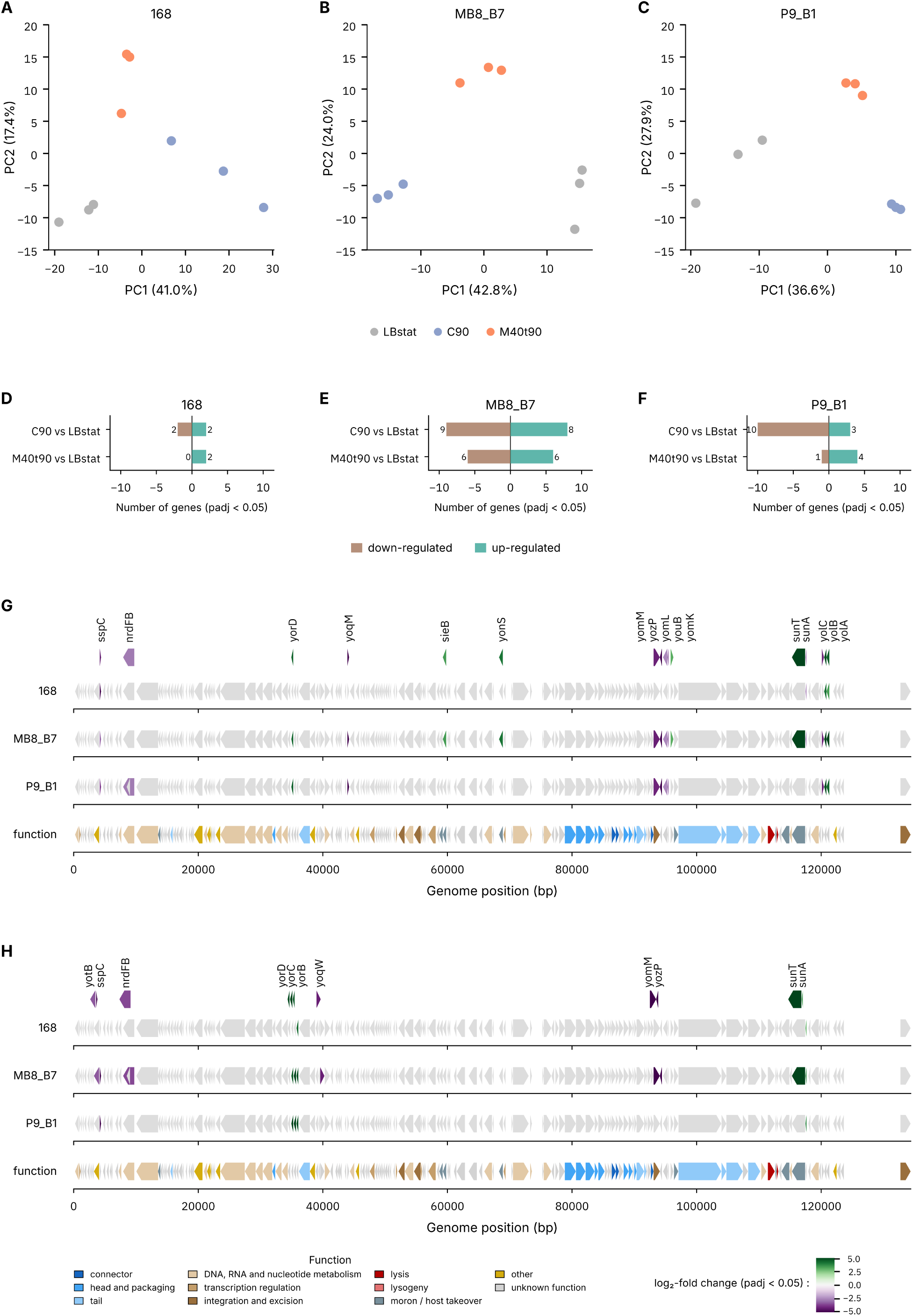
Prophage gene differential expression between conditions: C90 or M40t90 versus LBstat (lysogens only). PCA plots represent the difference between conditions for 168 (A), MB8_B7 (B) and P9_B1 (C) prophage. Number of differentially expressed prophage genes: C90 or M40t90 versus LBstat (lysogens only) (padj < 0.05) for (D) 168, (E) MB8_B7 and (F) P9_B1. (G) Gene expression pattern across the full SPβ genome for C90 versus LBstat (padj < 0.05). The bottom track represents the gene functions – derived from the *B. subtilis* 168 prophage annotation file. (H) Gene expression pattern across the full SPβ genome for M40t90 versus LBstat (padj < 0.05). The bottom track represents the gene functions – derived from the 168 prophage annotation file.

In the presence of mitomycin C, a stress-linked module was upregulated together with the sublancin precursor (*yorB*, *yorC*, *yorD*, *sunA* at padj < 0.05 and |log_2_FC| > 2) in MB8_B7 and P9_B1 and the sporulation-related gene *sspC* (at padj < 0.05 and |log_2_FC| > 2) was downregulated compared with LBstat (Figure S2 E and F). Additional genes were downregulated in MB8_B7 (*yozP*, *yoqW*, *yotB*, *yomM*) and two in P9_B1 (*yotM*, *yomM*). The core stress/sublancin upregulation was conserved across the three prophages (Figure 3 H and Figure S2 D, E and F). While the number of genes affected by the mitomycin C treatment was surprisingly low, MB8_B7 and P9_B1 lysSPβ growth rates were slower than their non-lysogen counterparts, with the exception of 168, which showed a decrease in growth rate after mitomycin C addition and, after 15 min, a growth rate similar to that of the two non-lysogens (Figure S3 B). In addition, we observed that the number of phage particles in culture (PFU/mL) with mitomycin C treatment increased by 1 (168, P9_B1) to 1.2 log_10_ (MB8_B7) relative to LBstat (Figure S3 H), indicating that phage particle production was triggered by the addition of mitomycin C to the medium compared with untreated cultures (Figure S3 G).

In competence medium versus LBstat, the response was broader (17 significant genes in MB8_B7, of which 6 were upregulated and 4 downregulated at |log_2_FC| > 2, and 13 significant genes in P9_B1, of which 3 were upregulated and 6 down-regulated at |log_2_FC| > 2). Among the upregulated genes were *yolA* and *yolB* (both unknown) and the stress-related gene *yorD*, conserved in both strains. Among the downregulated genes were the sporulation-related gene *sspC*, the sublancin precursor *sunA*, *yoqM* (unknown), *yolC* (SOS response), *yozP* (unknown), *yomM* (putative integrase), *yomL* (unknown), *youB* (unknown) and unnamed genes (CDS_0135, CDS_0127) (Figure 3 G; Figure S2 B and C). Only the upregulated *yolA* and *yolB* and the downregulated *sspC* and *sunA* were conserved in the 168 prophage (Figure 3 G; Figure S2 A).

As expected, the prophage response to different environmental conditions differed in the laboratory strain *B. subtilis* 168 and in the wild strains MB8_B7 and P9_B1. However, the pattern of upregulated and downregulated genes that we observed differed from the one predicted from the analysis of the public dataset.

### The prophage preserves the host transcriptional response to culture conditions but modulates the response magnitude

We next investigated whether the presence of a prophage influences the host’s transcriptional response to different culture conditions. We therefore examined whether lysogens and non-lysogens respond to a change in growth condition in the same way. The number of differentially regulated bacterial transcripts (padj < 0.05) was compared between C90 (or M40t90) and LBstat, based on their lysogeny status. Because of the large number of genes differentially expressed between culture conditions for both lysogen and non-lysogen strains (between 519 and 2,129 genes; Figure 4 E and F), we based the interpretation on Gene Ontology enrichment (Figure 4 A to D).

**Figure 4.**
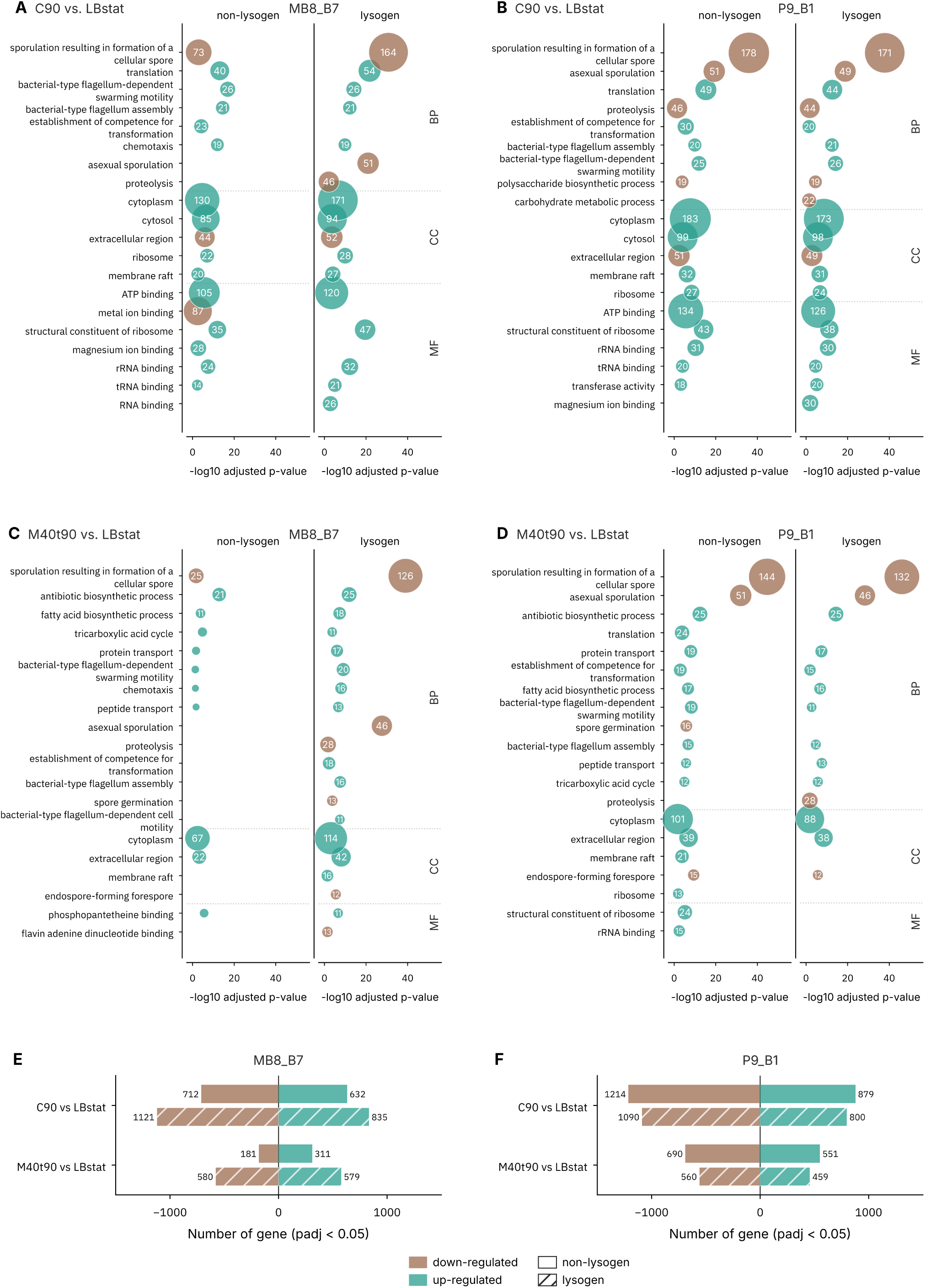
Gene Ontology enrichment on differentially expressed bacterial genes: C90 or M40t90 versus LBstat. Side-by-side comparison between non-lysogen (left facet) and lysogen (right facet) Gene Ontology enrichment for C90 vs. LBstat: MB8_B7 (A) and P9_B1 (B); and for M40t90 vs. LBstat: MB8_B7 (C) and P9_B1 (D). Positively enriched GO terms are in teal and negatively enriched terms are in brown. The top-20 terms per comparison are represented, ranked by p-value (padj < 0.05). Bubble area = number of significant genes. Terms are grouped by GO namespace: biological process (BP), cellular compartment (CC) and molecular function (MF) with dotted separators. Number of differentially expressed bacterial genes: C90 or M40t90 versus LBstat for (E) MB8_B7 and for (F) P9_B1 (padj < 0.05).

Regulation trends were shared between lysogens and non-lysogens (Figure 4 A to D). Sporulation genes were downregulated and motility (flagellum/swarming) genes were upregulated relative to LBstat under both mitomycin C and competence culture conditions, in all strains, regardless of prophage presence. Under competence, translation and ribosomal gene activity increased compared with LBstat, while amino-acid and nucleotide biosynthesis were downregulated(Figure 4 A and B). The response to mitomycin C treatment was less conserved between strains (Figure 4 C and D).

The amplitude of the transcriptional changes in different culture conditions was larger in the MB8_B7 lysogen than in its ΔSPβ counterpart (2,001 versus 1,348 genes in C90; 1,189 versus 519 in M40t90; Figure 4 E), while P9_B1 lysogen and non-lysogen yielded very similar numbers of condition-responsive genes (2,129 and 2,085 in C90; 1,240 and 1,281 in M40t90; Figure 4 F). We also observed that competence gene transcript enrichment was consistently higher for the non-lysogen strains, and conversely, downregulation of sporulation gene transcripts was stronger in the lysogen. The presence of the prophage largely did not alter the general trend of the host response to culture conditions; however, it modulated the response magnitude in a strain-dependent way.

### Prophage gene expression is largely independent of the host genetic background

We further asked whether prophage gene expression is influenced by host genetic background within a given culture condition. To make the three prophages comparable, we restricted the analysis to the 167 consensus genes present as one-to-one orthologues in all three prophage genomes (Table S4 and Figure S1).

There was no significant difference in the expression of consensus prophage genes between the two natural isolates P9_B1 and MB8_B7 in any of the three conditions (no gene at padj < 0.05; Figure 5 A, B, C). Compared with MB8_B7, the prophage in the laboratory strain 168 differed in only one gene (at padj < 0.05 and |log_2_FC| > 2; Figure 5 D): *yorC* was significantly downregulated in 168 compared with MB8_B7.

**Figure 5.**
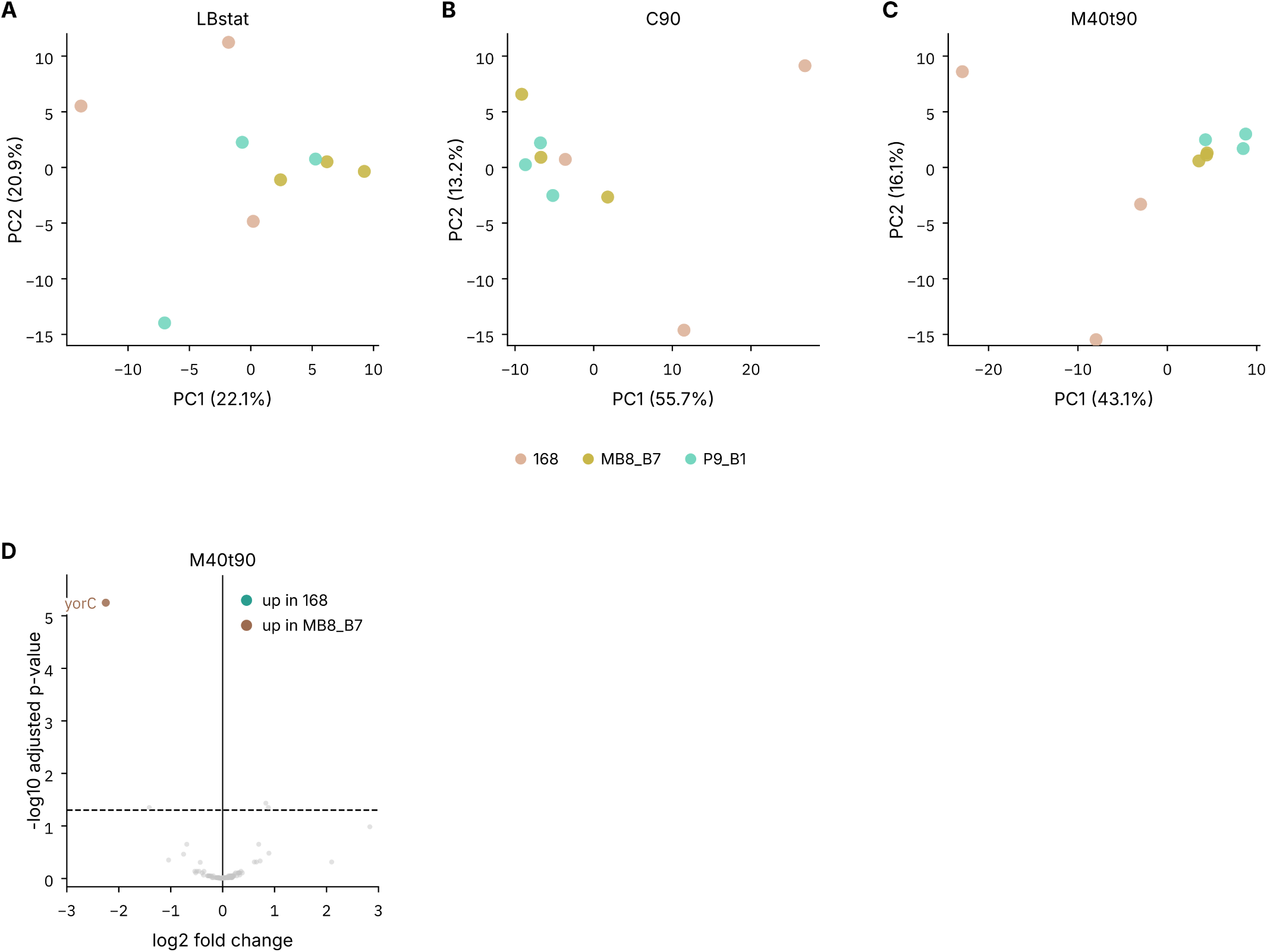
Differential gene expression comparison between the prophages of 168, MB8_B7 and P9_B1 across the three conditions. PCA for each condition: LBstat (A), C90 (B) and M40t90 (C). Red: 168, yellow: MB8_B7 and green: P9_B1. (D) Volcano plot for 168 versus MB8_B7 with upregulated genes in 168 shown in teal and in MB8_B7 in brown. Significance: at padj < 0.05 and |log_2_FC| > 2

Host background therefore had very little influence on the prophage transcriptional pattern in all culture conditions tested, with a minor deviation for strain 168.

### The prophage affects host transcription in a strain-dependent manner but consistently downregulates competence genes

We finally asked how the presence of the prophage affects host gene expression, and whether any general regulatory effects are shared by the two hosts. Whereas the previous analysis compared culture conditions within a strain (C90 or M40t90 versus LBstat), here we compared lysogens versus non-lysogens within the same culture condition. This was done separately for MB8_B7 and P9_B1, in each of the three conditions (LBstat, C90 and M40t90).

We observed that the differential expression pattern between lysogens and non-lysogens differed between the two strain backgrounds and that it was much stronger for MB8_B7 than for P9_B1 (Figure 6 A and B; PCA in Figure S5 A and B).

**Figure 6.**
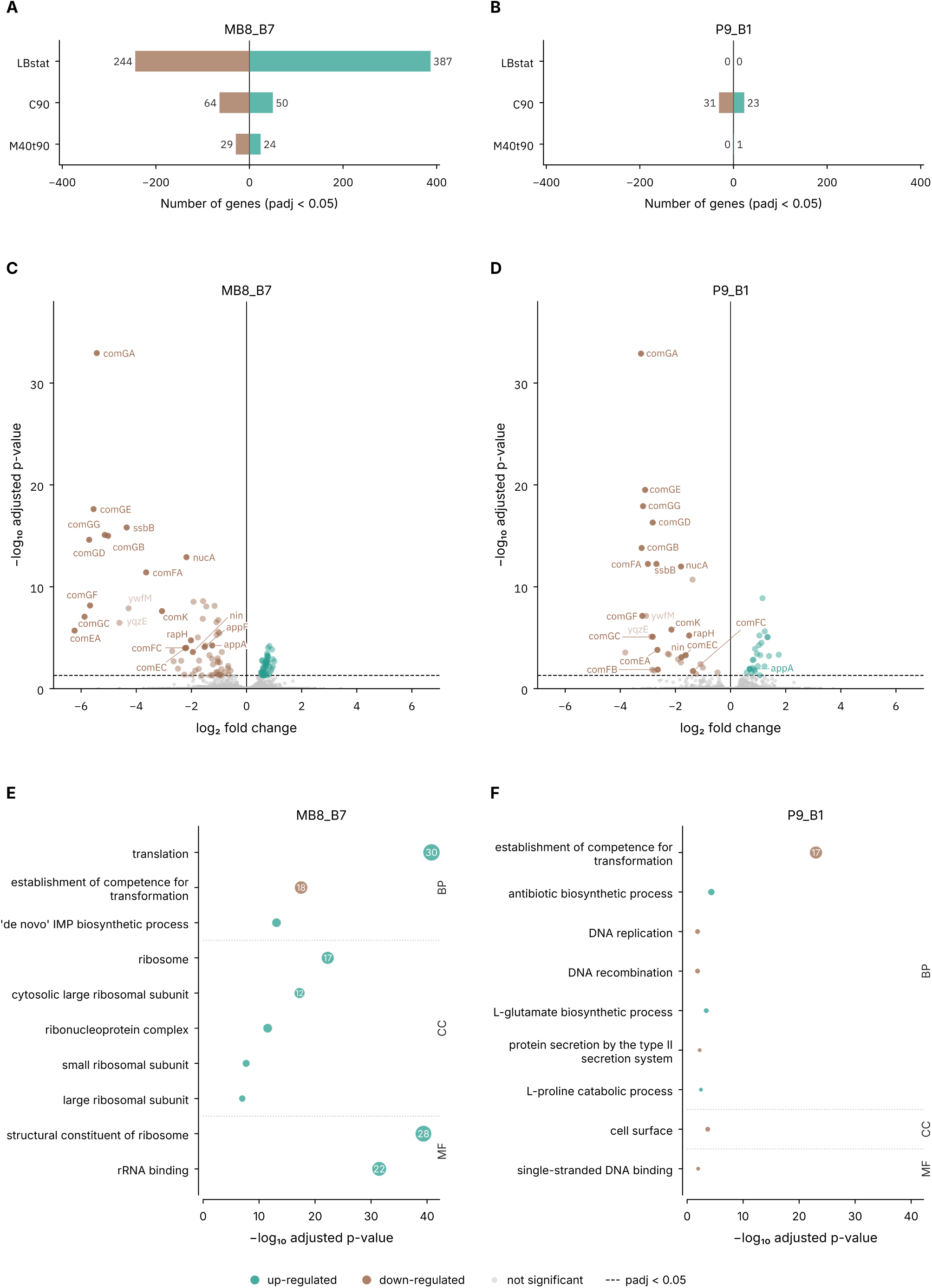
Differential expression between lysogen vs. non-lysogen (bacterial genes only) during growth in competence medium. Number of genes differentially expressed (padj < 0.05) for MB8_B7 (A) and P9_B1 (B) lysogen compared with the non-lysogen. Volcano plot showing differentially expressed genes between the lysogen and non-lysogen for MB8_B7 (C) and P9_B1 (D). Competence-related genes are highlighted on the plot. GO enrichment plots for MB8_B7 (E) and P9_B1 (F). The GO enrichment plots show the 12 most enriched terms per ontology ranked by p-value (padj < 0.05). Bubble area = number of significant genes. Terms are grouped by GO namespace: biological process (BP), cellular compartment (CC) and molecular function (MF) with dotted separators. Positively enriched genes/terms are shown in teal and negatively enriched terms in brown.

For the MB8_B7 strain, we observed significant changes between the lysogen and the non-lysogen strains under the different culture conditions (Figure 6 A), particularly in LBstat (Figure S4 A). The genes involved in sporulation were significantly upregulated in the lysogen and secreted proteins and catabolic processes such as the acetoin catabolic pathway were down-regulated (Figure S4 C and E). We also observed that bacterial concentration at sampling time was 0.5 log_10_ lower for the lysogen than for the non-lysogen although no difference was observed in biomass over time (Figure S3 A and D). In competence medium, transcription of competence genes was significantly decreased in the presence of the SPβ prophage (*comK*, *comGA/GB/GD/GE/GF/GG*, *comEA*, *comFA*, *ssbB*, *nucA* at padj < 0.05) and genes related to transcription and translation were upregulated (Figure 6 C and E). However, growth in competence medium and bacterial concentration at sampling time were similar for both lysogen and non-lysogen (Figure S3 C and F). Under mitomycin C treatment, we observed an activation of the catabolism of alternative nitrogen sources and the use of fatty acids as carbon sources through β-oxidation, and a decrease in central carbon metabolism, amino-acid biosynthesis, cell envelope maintenance and secondary metabolism such as acetoin; however, no GO term was noticeably enriched, unlike in the two other conditions (Figure S6 C and E).

The response of the P9_B1 strain differed from that of MB8_B7 for the three conditions. During growth in LBstat, there was no significant difference in expression between the lysogen and non-lysogen strains (no gene at padj < 0.05; Figure S4 B and D). In competence medium, as for the MB8_B7 strain, competence gene expression was decreased in the lysogen (54 genes, Figure 6 D and F). Under mitomycin C, lysogen and non-lysogen samples clustered separately for MB8_B7 but overlapped for P9_B1 (Figure S6 A and B), and only one gene, *yddT*, was significantly upregulated in the P9_B1 lysogen relative to the non-lysogen (Figure S6 D). This difference between the MB8_B7 and P9_B1 strains was also reflected in their growth and in the bacterial and phage particle concentrations (Figure S3).

Prophage presence thus influenced host gene expression, but the majority of the effects were host-dependent despite close genetic backgrounds. Nevertheless, we identified a consistent regulatory pattern across all tested host genetic backgrounds: the downregulation of competence-related genes. This observation suggests a conserved phage-host interaction during lysogeny, which we therefore selected for further investigation.

### SPβ reduces transformation efficiency by lowering the fraction of cells that enter the late competence state

To assess whether the difference in competence gene expression between lysogens and non-lysogens was reflected in the transformation efficiency of the strains, we further compared this efficiency between lysogen and non-lysogen strains.

For the P9_B1 strain, we observed that lysogen transformation frequency was consistently lower than for the non-lysogen strain for each type of DNA we tested (Figure 7 A, Table 3). It was significantly lower for gDNA *amyE*::P_pL_-*gfp*, with p-value < 0.01 (Table 3). We could not quantify transformation frequency in MB8_B7 due to the presence of *comI* (Smith et al., 2024) and the resulting low number of transformants.

**Figure 7.**
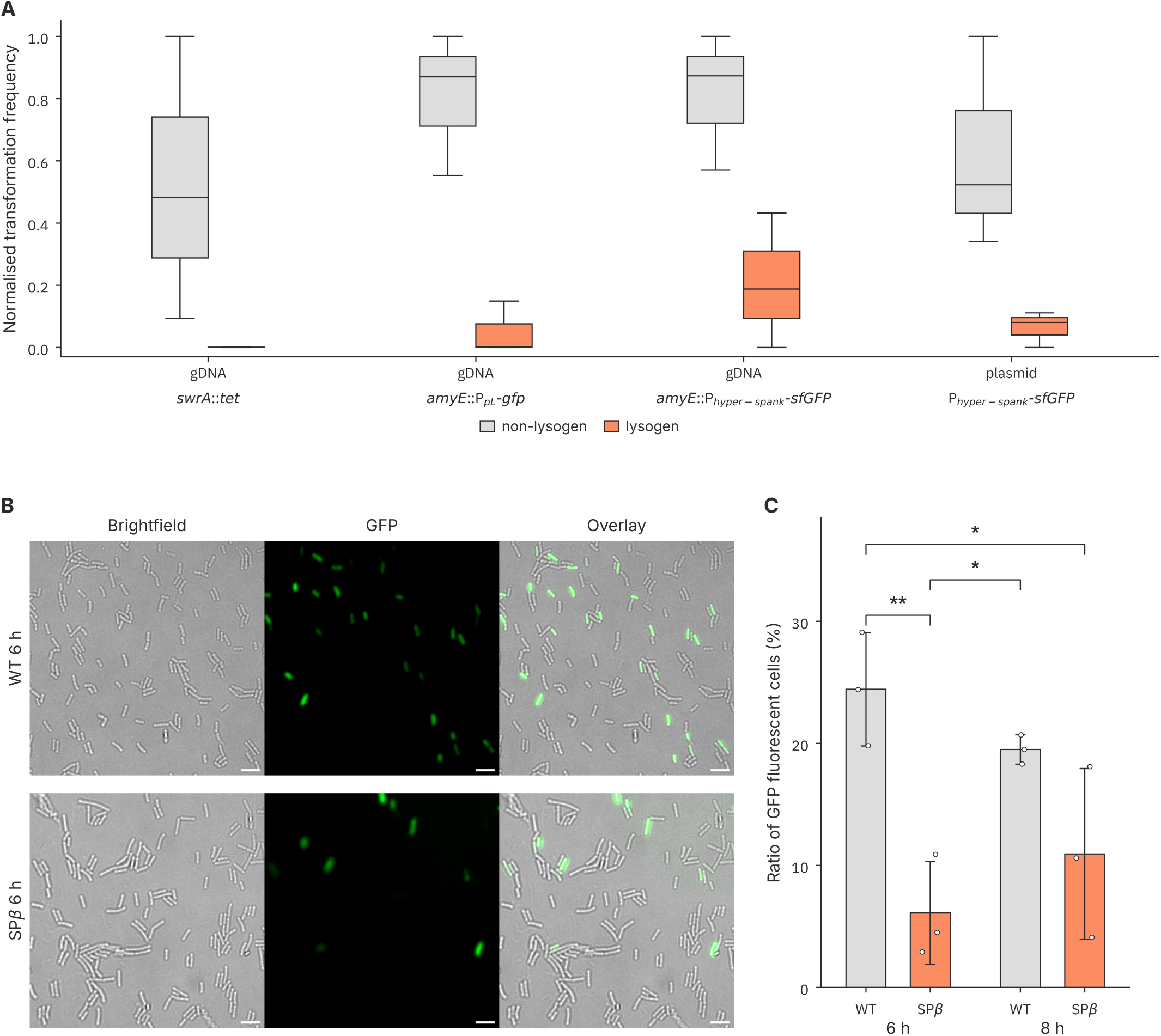
Bacterial cell competence in lysogens versus non-lysogens. (A) Min–max normalised transformation frequency for the different transformation assays. (B) Fraction of cells expressing P*_comGA_*-*gfp* at 6 h. Representative images of *B. subtilis* P9_B1 wild-type (top) and SPβ lysogen strain (bottom) carrying P*_comGA_*-*gfp*. Brightfield (left), ComGA-GFP fluorescence (middle) and overlay (right) are shown. Each panel shows a 62 × 62 µm region from a single tile of a stitched tilescan. Images were acquired with identical exposure and laser settings, and the GFP channel is displayed with identical linear contrast in all panels. Scale bars, 5 µm. (C) Percentage of P*_comGA_*-*gfp*-positive cells in WT and SPβ cultures at 6 h and 8 h. Bars show the mean ± SD of three biological replicates. Cells analysed per condition: WT 6 h, n = 3,697; SPβ 6 h, n = 2,422; WT 8 h, n = 3,100; SPβ 8 h, n = 600. Statistical significance was determined by one-way ANOVA followed by Tukey’s post hoc test (*p < 0.05, **p < 0.01); all other comparisons not significant. Non-lysogens are shown in grey and lysogens in orange.

**Table 3.**
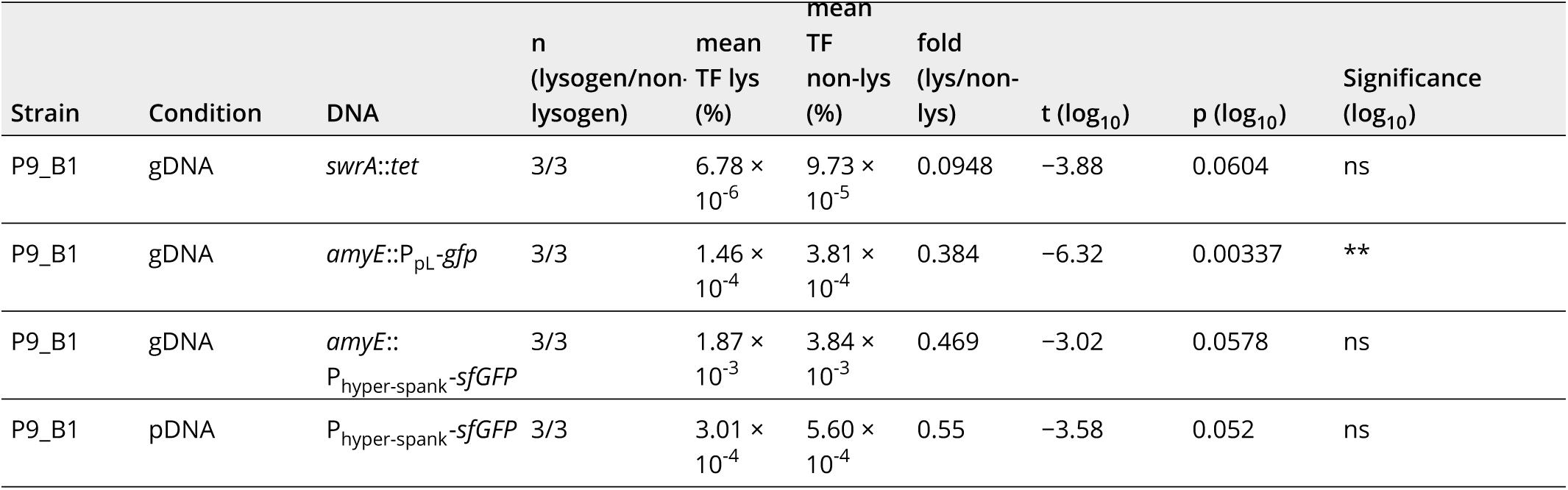
Transformation frequency in P9_B1 strains. Statistical significance between lysogens and non-lysogens was determined using a two-sided Welch’s t-test on log_10_-transformed transformation frequency for each experiment.

| Strain | Condition | DNA | n (lysogen/non-lysogen) | mean TF lys (%) | mean TF non-lys (%) | fold (lys/non-lys) | t (log<sub>10</sub>) | p (log<sub>10</sub>) | Significance (log<sub>10</sub>) |
|---|---|---|---|---|---|---|---|---|---|
| P9_B1 | gDNA | *swrA*::*tet* | 3/3 | 6.78 × 10<sup>-6</sup> | 9.73 × 10<sup>-5</sup> | 0.0948 | −3.88 | 0.0604 | ns |
| P9_B1 | gDNA | *amyE*::P<sub>pL</sub>-*gfp* | 3/3 | 1.46 × 10<sup>-4</sup> | 3.81 × 10<sup>-4</sup> | 0.384 | −6.32 | 0.00337 | ** |
| P9_B1 | gDNA | *amyE*::P<sub>hyper-spank</sub>-*sfGFP* | 3/3 | 1.87 × 10<sup>-3</sup> | 3.84 × 10<sup>-3</sup> | 0.469 | −3.02 | 0.0578 | ns |
| P9_B1 | pDNA | P<sub>hyper-spank</sub>-*sfGFP* | 3/3 | 3.01 × 10<sup>-4</sup> | 5.60 × 10<sup>-4</sup> | 0.55 | −3.58 | 0.052 | ns |

We also monitored the activation of the late competence genes, genes controlled by the P*_comGA_* promoter. At 6 h, the percentage of non-lysogenic cells entering the late competence stage was higher than in the lysogen, as we observed more fluorescent cells for the non-lysogens (24 % versus 6 %; p-value < 0.01) (Figure 7 B and C). At 8 h, the number of cells in the competent state decreased for the wild-type, but the proportion of fluorescent cells was still higher than for the lysogen (n.s.).

Altogether, these results are consistent with the RNA-seq observation: the prophage negatively affected the transformation rate and fewer cells were in the late competence state when the prophage was present.

## Discussion

Large transcriptomic datasets may prove to be a valuable source of information about prophage-host interaction. By analysing an existing transcriptomic dataset (Nicolas et al., 2012), we were able to identify suitable culture conditions in which to study prophage-host interaction. Natural competence appeared to be a particularly interesting condition: gene expression variation pattern is very similar to that of mitomycin C, a trigger of the SPβ lytic cycle (Warner et al., 1977; Kohm and Hertel, 2021), despite the fact that SPβ is known not to enter the lytic cycle during competence (McVeigh and Yasbin, 1996).

We showed that competence genes were downregulated in lysogens compared with non-lysogens, and this effect was reproducible in the two *B. subtilis* natural isolates tested, isolates with different genetic backgrounds. It is known that mobile genetic elements (MGEs), in general, and prophages, in particular, can suppress bacterial transformation to prevent chromosome curing, using different strategies (Kwun et al., 2025). One common strategy is for the prophage to lyse the host cells when the host enters competence as shown for phi105 and SPO2 (Garro and Law, 1974; McVeigh and Yasbin, 1996), which would explain the decrease in competence gene transcripts. However, there was no difference in bacterial concentration between lysogens and non-lysogens when cultivated in competence medium, suggesting the absence of cell lysis and therefore another mechanism for competence gene regulation. Phage particle concentrations measured after cultivation in competence medium or culture in LB during stationary phase were comparable for equivalent bacterial concentrations, indicating no increase in phage particle production and corroborating this result and the previous conclusion of McVeigh and Yasbin (1996) on phi3T, another prophage from the *Spbetavirus* genus. Another tactic promoted by MGEs is to down-regulate competence gene expression or block the competence apparatus. This strategy has been mostly observed for plasmids (Vesel et al., 2025), such as pLS20 (Singh et al., 2012), pBS32 (Konkol et al., 2013), pLS32 (Burton and Kearns, 2020); prophages seem to favour integration in or near the competence genes (Vesel et al., 2025). However, SPβ is integrated in the *spsM* gene and all known *Spbetavirus* are integrated in the same area of the genome (within 1 % of the genome length of one another (Kohm and Hertel, 2021; Kohm et al., 2022)), distant from the competence gene cluster. Our results clearly suggest that SPβ affects the regulation of the competence genes, even though the earlier study by McVeigh and Yasbin (1996) states that transformation is not affected by the close relative of SPβ, phi3T. However, they used transformation frequency as the readout and we also observed through this study that depending on the form of transforming DNA used and medium batch, transformation frequency may not appear significantly affected even though it tends to be lower for the lysogen. Fluorescence microscopy imaging showed a difference in the proportion of cells entering the late competence stage between the lysogen and non-lysogen, supporting our transcriptomic data. In our study, two prophage genes were significantly upregulated during culture in competence medium: *yolA* coding for a secreted protein (Antelmann et al., 2001; Tjalsma et al., 2004) and *yolB*, for which we could not find evidence of their function or mechanism of action in the literature. Based on our results, we hypothesise that the two SPβ genes are important for either downregulating competence genes or maintaining the prophage state during competence.

Our results also indicate that the presence of the prophage does not completely alter its host transcriptomic pattern. Rather, it modulates some aspects to its advantage. Competence genes are still upregulated in competence medium even though they are negatively modulated by the prophage as previously described. The same applies during growth in LB medium to stationary phase, when the availability of nutrients in the medium decreases. All strains showed an increase in sporulation gene expression (Piggot and Hilbert, 2004). In *B. subtilis* MB8_B7, the presence of the prophage seems to lead to the upregulation of sporulation-related transcripts, which we interpret as a preservation mechanism. The more spores are formed, the more phages can survive the period of nutrient scarcity in the lysogenic state and would be able to recolonise their environment (Butala and Dragoš, 2023) when conditions are more favourable for the bacteria to thrive. However, we also noticed a significant decrease in acetoin catabolism, although production and consumption of acetoin have been shown to fuel the cell in the absence of glucose and during spore formation (Yoshida et al., 2000; Ould Ali et al., 2001).

Interestingly, there was no noticeable transcriptomic difference between *B. subtilis* P9_B1 lysogen and non-lysogen in stationary phase or when cultivated in the presence of mitomycin C. It appears that the presence of the prophage does not influence its host under those culture conditions. Thus, the presence of the SPβ prophage seems to affect each of the strains differently. According to our observations, the difference could not be attributed to host-dependent changes in prophage regulation, as no difference in prophage gene expression pattern was observed when comparing prophages residing in different hosts cultivated under the same conditions. Although the SPβ prophage used to lysogenise *B. subtilis* P9_B1 was induced from MB8_B7 (Dragoš et al., 2021a) to allow for comparison, the analysis of the genome revealed that seven genes were missing compared with the original prophage genome. Thus, we cannot exclude the possibility that one or more of the seven missing genes (including a nuclease and six hypothetical proteins) might account for this difference. Alternatively, this could indicate that each of the hosts responds differently to prophage presence. *B. subtilis* MB8_B7 is a natural lysogen for SPβ (Kiesewalter et al., 2021) in contrast to *B. subtilis* P9_B1 (Kiesewalter et al., 2021), a natural non-lysogen for SPβ, which was lysogenised in the laboratory (Dragoš et al., 2021a). We also reflected on whether there is any difference between the genetic backgrounds of the two strains that might explain why the prophage would be more adapted to one strain than to the other and maybe was conserved in MB8_B7 but not in P9_B1 even though the phage is able to infect and lysogenise the strain. It has been shown that despite the ability of prophages to infect a strain, in natural settings, the strains maintain prophages depending on prophage-encoded benefits and fitness factors (Dragoš et al., 2021a) and host regulatory elements required for maintenance of the prophage such as SigY for SPβ (Mendez et al., 2012). In order to test whether the difference is dependent on strain genomic background or linked to the evolutionary history of lysogeny, we would have to perform transcriptomic analyses on multiple strains isolated with and without the SPβ prophage and compare the results.

Surprisingly, we observed few transcriptomic changes in the prophage transcriptome under mitomycin C treatment. Mitomycin C is a general SOS inducer (Baharoglu and Mazel, 2014) and is routinely used in laboratories to induce prophages. SPβ prophage is inducible with mitomycin C (Warner et al., 1977; Kohm and Hertel, 2021) and its activation is RecA/LexA dependent and part of the SOS response (Goranov et al., 2006). Only three genes stand out on the prophage side: *yorB*, *yorC* and *yorD*, a stress protein (Salzberg et al., 2015). Those three genes are known to be possibly activated by the SOS response as an SOS box has been identified upstream of *yorB* (Lazarevic et al., 1997; Au et al., 2005) even though the function and mechanism are unknown (Kohm and Hertel, 2021). Their activation is consistent with DNA damage SOS response activation in bacteria in presence of mitomycin. However, we were expecting to see upregulation of the prophage transcription machinery and structural proteins (Kohm et al., 2022, 2023) and of the recombination directionality factor *sprB* (Abe et al., 2014). The result could be explained by the concentration of 40 ng/mL mitomycin C used in this study, which is lower than the optimal concentration of 1 µg/mL for SPβ induction according to Warner et al. (1977) and for which *B. subtilis* strain can still grow according to Hoch et al. (1967). Our own data confirmed that the effect of the mitomycin C on the lysogen was mild; it only showed a slower growth rate than the non-lysogen and only 1- to 2-log (for MB8_B7 lysogen) difference in phage particle production compared with the stationary phase condition. We therefore attribute the weak response to the low mitomycin C concentration rather than to a problem with the RNA-seq data. The presence and maintenance of a large lysogen population likely contribute to a high AimP (arbitrium peptide) concentration and the consequent repression of the arbitrium system, reducing the percentage of cells entering the lytic cycle (Chmielowska et al., 2026). Still, public dataset analysis obtained from laboratory strain *B. subtilis* 168 (Nicolas et al., 2012) revealed that even at the low mitomycin C concentration, the regulatory genes, structural proteins and recombination directionality factor transcription are upregulated compare with the other conditions, therefore we assume that difference is likely due to not being able to reproduce the exact condition as in the original study. Another possibility could involve SPβ interaction with ICEBs1 (Johnson et al., 2022; McKeithen-Mead et al., 2025; Cai et al., 2026), present in 168 and absent from our strains (Stefanic et al., 2025).

This work establishes a workflow harnessing a public dataset to discover new mechanisms of host control by phages. It indicates that prophage gene expression is independent of the host genetic background whereas the phage effect on the host is influenced by the host genetic background. This work identifies competence as a condition in which the prophage affects its host and our experimental data confirmed a downregulation of the competence genes and a reduction of the competent cell population. Further investigation of this mechanism could contribute to a better understanding of SPβ prophage interaction with its host and better predict the outcome of transformation in lysogenised *B. subtilis* strains. Moreover, a similar approach could be extended to other prophage-rich model bacteria, leveraging existing transcriptomic datasets to systematically investigate phage-host interactions and prioritise the most relevant culture conditions for experimental exploration.

## Data availability

The engineered bacterial strains created for this study are available from the corresponding author upon reasonable request. The raw transcriptomic sequencing data (RNA-seq) generated during this study have been deposited in the European Nucleotide Archive (ENA) under the project accession number PRJEB129176.

## Acknowledgements

This work was supported by the European Research Council (ERC) under the European Union’s Horizon Europe research and innovation programme (ERC Starting Grant PHAGECONTROL, grant agreement no. [101041421] to A.D.). V.G. was supported by PHAGECONTROL, Views and opinions expressed are however those of the author(s) only and do not necessarily reflect those of the European Union or the European Research Council Executive Agency. Neither the European Union nor the granting authority can be held responsible for them. A.D. and V.G. were also supported by the Slovenian Agency for Research and Innovation [Grant number P4-0116]. L.G. and J.U. were supported by the Slovenian Agency for Research and Innovation [Grant number N1-0426]. Š.B. was supported by the Slovenian Agency for Research and Innovation [Grant Nos. P4-0463, J4-50134 and J7-70268]. We thank Sara Fišer for technical support. We also thank Tajda Klobučar and Urška Janjoš for technical support with the RNA-seq experiments, and Klara Kuret Hodnik for advice on the bioinformatic analyses.

## Author contributions

V.G. and A.D. designed the study. V.G. performed the vast majority of the experiments, with the help of A.D. and L.G. Š.B. largely contributed to RNA extraction optimisation. L.G. and J.U. largely contributed to RNA library preparation, QC and data analysis, V.G. and A.D. wrote the manuscript. All authors contributed to the final version.

