## Supplemental Materials for "Competence repression by the SPβ prophage in *Bacillus subtilis* revealed by transcriptomics across culture conditions"

### Supplementary Data

**Supplementary Results S1** Dataset structure and quality assessment.

Among the 102 conditions in the study, only three have technical replicates. Conditions with exactly three biological replicates form the largest group; other conditions have between one and six biological replicates (the maximum of six is observed for condition S6). There are 17 conditions with two biological replicates and 17 with only a single replicate. The latter mainly concern batch fermentation in modified M9 medium and were excluded from condition selection and statistical analyses. Replicate MG+120_2 was also excluded from the analysis because the condition name (120) and the condition description (150) were contradictory. In addition, the preliminary data quality analysis informed us that some GeneSymbol names were repeated across IDs as *yoyK* for the prophage genes (ID 3078 - ORF BSU21598 and ID 3079 - ORF BSU21599). However, IDs and ORFs were all unique and could be used to identify the target (further in the text simplified to gene as most targets are designed on individual genes).

Despite the low number of biological replicates (only 2 for 16 of the conditions), changes in expression were considered biologically meaningful because within-condition dispersion was not significant.

**Table S1** Samples collected during the study for RNA-seq analysis. The table contains the sample ID, the corresponding strain and genotype, the biological replicate identifier, whether the strain carries the SPβ prophage and the indexed primer pair used for library preparation.

| ID | Condition | Strain | Genotype | Biological replicate | Has prophage | Indexed primer IDs |
|---|---|---|---|---|---|---|
| 16 | LBstat | 168 | wt | 4 | True | 501/701 |
| 17 | LBstat | P9_B1 | wt | 4 | False | 501/702 |
| 18 | LBstat | P9_B1 | SPbeta | 4 | True | 501/703 |
| 19 | LBstat | MB8_B7 | delta_SPbeta | 4 | False | 501/704 |
| 20 | LBstat | MB8_B7 | wt | 4 | True | 501/705 |
| 21 | LBstat | 168 | wt | 5 | True | 501/706 |
| 22 | LBstat | P9_B1 | wt | 5 | False | 501/707 |
| 23 | LBstat | P9_B1 | SPbeta | 5 | True | 501/708 |
| 24 | LBstat | MB8_B7 | delta_SPbeta | 5 | False | 502/701 |
| 25 | LBstat | MB8_B7 | wt | 5 | True | 502/702 |
| 26 | LBstat | 168 | wt | 6 | True | 502/703 |
| 27 | LBstat | P9_B1 | wt | 6 | False | 502/704 |
| 28 | LBstat | P9_B1 | SPbeta | 6 | True | 502/705 |
| 29 | LBstat | MB8_B7 | delta_SPbeta | 6 | False | 502/706 |
| 30 | LBstat | MB8_B7 | wt | 6 | True | 502/707 |
| 31 | M40t90 | 168 | wt | 1 | True | 502/708 |
| 32 | M40t90 | P9_B1 | wt | 1 | False | 503/701 |
| 33 | M40t90 | P9_B1 | SPbeta | 1 | True | 503/702 |
| 34 | M40t90 | MB8_B7 | delta_SPbeta | 1 | False | 503/703 |
| 35 | M40t90 | MB8_B7 | wt | 1 | True | 503/704 |
| 36 | M40t90 | 168 | wt | 2 | True | 503/705 |
| 37 | M40t90 | P9_B1 | wt | 2 | False | 503/706 |
| 38 | M40t90 | P9_B1 | SPbeta | 2 | True | 503/707 |
| 39 | M40t90 | MB8_B7 | delta_SPbeta | 2 | False | 503/708 |
| 40 | M40t90 | MB8_B7 | wt | 2 | True | 504/701 |
| 41 | M40t90 | 168 | wt | 3 | True | 504/702 |
| 42 | M40t90 | P9_B1 | wt | 3 | False | 504/703 |
| 43 | M40t90 | P9_B1 | SPbeta | 3 | True | 504/704 |
| 44 | M40t90 | MB8_B7 | delta_SPbeta | 3 | False | 504/705 |
| 45 | M40t90 | MB8_B7 | wt | 3 | True | 504/706 |
| 46 | C90 | 168 | wt | 1 | True | 504/707 |
| 47 | C90 | P9_B1 | wt | 1 | False | 504/708 |
| 48 | C90 | P9_B1 | SPbeta | 1 | True | 505/701 |
| 49 | C90 | MB8_B7 | delta_SPbeta | 1 | False | 505/702 |
| 50 | C90 | MB8_B7 | wt | 1 | True | 505/703 |
| 51 | C90 | 168 | wt | 2 | True | 505/704 |
| 52 | C90 | P9_B1 | wt | 2 | False | 505/705 |
| 53 | C90 | P9_B1 | SPbeta | 2 | True | 505/706 |
| 54 | C90 | MB8_B7 | delta_SPbeta | 2 | False | 505/707 |
| 55 | C90 | MB8_B7 | wt | 2 | True | 505/708 |

| ID | Condition | Strain | Genotype | Biological replicate | Has prophage | Indexed primer IDs |
|---|---|---|---|---|---|---|
| 56 | C90 | 168 | wt | 3 | True | 506/701 |
| 57 | C90 | P9_B1 | wt | 3 | False | 506/702 |
| 58 | C90 | P9_B1 | SPbeta | 3 | True | 506/703 |
| 59 | C90 | MB8_B7 | delta_SPbeta | 3 | False | 506/704 |
| 60 | C90 | MB8_B7 | wt | 3 | True | 506/705 |

**Table S2** Primers used for the sequencing libraries. Spaces separate the constant, index and UMI segments of each sequence.

| Primer name | Sequence |
|---|---|
| randomer_RT | TCAGACGTGTGCTCTTCCGATCT NNNNNNN |
| tso_rGrGpN | /5Biosg/CACGACGCTCrUrUCCGATrCTNNNNGAGTNNNrGrG+N |
| i501 | AATGATACGGCGACCACCGAGATCTACAC CACCATTG ACACTCTTTCCCTACACGACGCTCTTCCGATCT |
| i502 | AATGATACGGCGACCACCGAGATCTACAC TTATCAGC ACACTCTTTCCCTACACGACGCTCTTCCGATCT |
| i503 | AATGATACGGCGACCACCGAGATCTACAC CATATGGT ACACTCTTTCCCTACACGACGCTCTTCCGATCT |
| i504 | AATGATACGGCGACCACCGAGATCTACAC TGCTAGAC ACACTCTTTCCCTACACGACGCTCTTCCGATCT |
| i505 | AATGATACGGCGACCACCGAGATCTACAC AGAAGTCA ACACTCTTTCCCTACACGACGCTCTTCCGATCT |
| i506 | AATGATACGGCGACCACCGAGATCTACAC GCTTCGGT ACACTCTTTCCCTACACGACGCTCTTCCGATCT |
| i701 | CAAGCAGAAGACGGCATACGAGAT TACCTAGC GTGACTGGAGTTCAGACGTGTGCTCTTCCGATC |
| i702 | CAAGCAGAAGACGGCATACGAGAT GATTCTCA GTGACTGGAGTTCAGACGTGTGCTCTTCCGATC |
| i703 | CAAGCAGAAGACGGCATACGAGAT ATAGGAAG GTGACTGGAGTTCAGACGTGTGCTCTTCCGATC |
| i704 | CAAGCAGAAGACGGCATACGAGAT CTGCACCT GTGACTGGAGTTCAGACGTGTGCTCTTCCGATC |
| i705 | CAAGCAGAAGACGGCATACGAGAT GACGTGAG GTGACTGGAGTTCAGACGTGTGCTCTTCCGATC |
| i706 | CAAGCAGAAGACGGCATACGAGAT AGCTCTCC GTGACTGGAGTTCAGACGTGTGCTCTTCCGATC |
| i707 | CAAGCAGAAGACGGCATACGAGAT TTCTATGG GTGACTGGAGTTCAGACGTGTGCTCTTCCGATC |
| i708 | CAAGCAGAAGACGGCATACGAGAT GCACGGTA GTGACTGGAGTTCAGACGTGTGCTCTTCCGATC |
| P5_multiplex_primer | AATGATACGGCGACCACC |
| P7_multiplex_primer | CAAGCAGAAGACGGCATA |

**Table S3** Oligomers designed with Ribocutter (Wilkins and Ule, 2021) and used to generate the sgRNAs for rRNA depletion in the samples.

| Oligo name | Sequence |
|---|---|
| oVG47 | TTCTAATACGACTCACTATAGCTCAACCGGGGAGGGTCATGTTTTAGAGCTAGA |
| oVG48 | TTCTAATACGACTCACTATAGAGCCCCCGGCTCAACCGGGGGTTTTAGAGCTAGA |
| oVG49 | TTCTAATACGACTCACTATAGCCCCCGGCTCAACCGGGGAGTTTTAGAGCTAGA |
| oVG50 | TTCTAATACGACTCACTATAGAAGAGGCAAGCCTCTTGGTTGTTTTAGAGCTAGA |
| oVG51 | TTCTAATACGACTCACTATAGACGCATTTCACCGCTACACGGTTTTAGAGCTAGA |
| oVG52 | TTCTAATACGACTCACTATAGGGTCGGAACTTACCCGACAGTTTTAGAGCTAGA |
| oVG53 | TTCTAATACGACTCACTATAGCACGAAAGGCGCAACGATCTGTTTTAGAGCTAGA |
| oVG54 | TTCTAATACGACTCACTATAGGGGAGGGTCATTGGAAACTGTTTTAGAGCTAGA |
| oVG55 | TTCTAATACGACTCACTATAGCATTTGCTTTCTCTTCCTCCGTTTTAGAGCTAGA |
| oVG56 | TTCTAATACGACTCACTATAGAGAGGCAAGCCTCTTGGTTTGTTTTAGAGCTAGA |
| oVG57 | TTCTAATACGACTCACTATAGACACGACGCTCTTCCGATCTGTTTTAGAGCTAGA |
| oVG58 | TTCTAATACGACTCACTATAGTGAAAGCCCCCGGCTCAACCGTTTTAGAGCTAGA |
| oVG59 | TTCTAATACGACTCACTATAGAAGTCTGATGTGAAAGCCCCGTTTTAGAGCTAGA |
| oVG60 | TTCTAATACGACTCACTATAGCAGAACGTCGTGAGACAGTTGTTTTAGAGCTAGA |
| oVG61 | TTCTAATACGACTCACTATAGACCCCAAGAGGCAAGCCTCTGTTTTAGAGCTAGA |
| oVG62 | TTCTAATACGACTCACTATAGTAGGGAAAGAGTGTCAATGGGTTTTAGAGCTAGA |
| oVG63 | TTCTAATACGACTCACTATAGTATAGTACCTGTGAAGATGCGTTTTAGAGCTAGA |
| oVG64 | TTCTAATACGACTCACTATAGGATACCACTCCATTACGAGGTTTTAGAGCTAGA |
| oVG65 | TTCTAATACGACTCACTATAGAATGGTGGTGTAGATCTCGGGTTTTAGAGCTAGA |
| oVG66 | TTCTAATACGACTCACTATAGTGAAACATCTAAGTACCCGGGTTTTAGAGCTAGA |
| oVG67 | TTCTAATACGACTCACTATAGTTGCCACCTACGTATTACCGGTTTTAGAGCTAGA |
| oVG68 | TTCTAATACGACTCACTATAGACCACTCCATTACGAGTGGTGTTTTAGAGCTAGA |
| oVG69 | TTCTAATACGACTCACTATAGCTGATAAGTGTAGATCTCGGGTTTTAGAGCTAGA |
| oVG70 | TTCTAATACGACTCACTATAGAACTGAAACATCTAAGTACCGTTTTAGAGCTAGA |
| oVG71 | TTCTAATACGACTCACTATAGCGTTTCGCTCGCCGCTACTCGTTTTAGAGCTAGA |
| oVG72 | TTCTAATACGACTCACTATAGATCGGAAGAGCGTCGTGTAGTTTTAGAGCTAGA |
| oVG73 | TTCTAATACGACTCACTATAGATATCCTATGAATTCAGATAGTTTTAGAGCTAGA |
| oVG74 | TTCTAATACGACTCACTATAGTATCCATATCTGAATTCATGTTTTAGAGCTAGA |
| oVG75 | TTCTAATACGACTCACTATAGTTTCCCCATTCGGAAATCTCGTTTTAGAGCTAGA |
| oVG76 | TTCTAATACGACTCACTATAGACCCTCCCCGGTTGAGCCGGTTTTAGAGCTAGA |
| oVG77 | TTCTAATACGACTCACTATAGGGAGGGTCATTGGAAACTGGTTTTAGAGCTAGA |
| oVG78 | TTCTAATACGACTCACTATAGCAAGCGTTGTCCGGAATTATGTTTTAGAGCTAGA |
| oVG79 | TTCTAATACGACTCACTATAGAAACCAAGAGGCTTGCCTCTGTTTTAGAGCTAGA |
| oVG80 | TTCTAATACGACTCACTATAGCCCTTTACGCCCAATAATTCGTTTTAGAGCTAGA |
| oVG81 | TTCTAATACGACTCACTATAGGGCGTAAAGGGCTCGCAGGGTTTTAGAGCTAGA |
| oVG82 | TTCTAATACGACTCACTATAGTACCACTCCATTACGAGTGGGTTTTAGAGCTAGA |
| oVG83 | TTCTAATACGACTCACTATAGACCCACCACTCGTAATGGAGGTTTTAGAGCTAGA |
| oVG84 | TTCTAATACGACTCACTATAGTTGAGTGCAGAAGAGGAGAGGTTTTAGAGCTAGA |
| oVG85 | TTCTAATACGACTCACTATAGATTGGGCGTAAAGGGCTCGCGTTTTAGAGCTAGA |
| oVG86 | TTCTAATACGACTCACTATAGCAGTTTCCAATGACCCTCCCGTTTTAGAGCTAGA |

| Oligo name | Sequence |
|---|---|
| oVG87 | TTCTAATACGACTCACTATAGAGGTAGCGAAATTCCTTGTCGTTTTAGAGCTAGA |
| oVG88 | TTCTAATACGACTCACTATAGTGAAAGCCCCCGGCTCAACGTTTTAGAGCTAGA |
| oVG89 | TTCTAATACGACTCACTATAGACCCTCCCCGGTTGAGCCGGGTTTTAGAGCTAGA |
| oVG90 | TTCTAATACGACTCACTATAGGCTTGCCTCTTGGGGTTGTGTTTTAGAGCTAGA |
| oVG91 | TTCTAATACGACTCACTATAGAGTGTCCTACAACCCCAAGGTTTTAGAGCTAGA |
| oVG92 | TTCTAATACGACTCACTATAGTGACCCTCCCCGGTTGAGCCGTTTTAGAGCTAGA |
| oVG93 | TTCTAATACGACTCACTATAGCTGAGTAGCGGCGAGCGAAAGTTTTAGAGCTAGA |
| oVG94 | TTCTAATACGACTCACTATAGCGGGGAGGGTCATTGGAAACGTTTTAGAGCTAGA |
| oVG95 | TTCTAATACGACTCACTATAGCGGGATTAGCCCAAACCAAGGTTTTAGAGCTAGA |
| oVG96 | TTCTAATACGACTCACTATAGTCCGGAATTATTGGGCGTAAGTTTTAGAGCTAGA |
| oVG97 | TTCTAATACGACTCACTATAGTGAGTAGCGGCGAGCGAAACGTTTTAGAGCTAGA |
| oVG98 | TTCTAATACGACTCACTATAGGGAAACCCACCACTCGTAAGTTTTAGAGCTAGA |
| oVG99 | TTCTAATACGACTCACTATAGAGTGGAATTCCACGTGTAGGTTTTAGAGCTAGA |
| oVG100 | TTCTAATACGACTCACTATAGGTCGCCGTATCATTAAAAAGTTTTAGAGCTAGA |
| oVG101 | TTCTAATACGACTCACTATAGCGTAGGTGGCAAGCGTTGTCGTTTTAGAGCTAGA |
| oVG102 | TTCTAATACGACTCACTATAGTCAATGGTGGTGTAGATCTGTTTTAGAGCTAGA |
| oVG103 | TTCTAATACGACTCACTATAGAGATCGGAAGAGCGTCGTGTGTTTTAGAGCTAGA |
| oVG104 | TTCTAATACGACTCACTATAGAAGCGTTGTCCGGAATTATTGTTTTAGAGCTAGA |
| oVG105 | TTCTAATACGACTCACTATAGAAAGCCCCCGGCTCAACCGGTTTTAGAGCTAGA |
| oVG106 | TTCTAATACGACTCACTATAGAAATGCGATTCCCTGAGTAGGTTTTAGAGCTAGA |
| oVG107 | TTCTAATACGACTCACTATAGACCAAGAGGCTTGCCTCTTGGTTTTAGAGCTAGA |
| oVG108 | TTCTAATACGACTCACTATAGATGACCCTCCCCGGTTGAGCGTTTTAGAGCTAGA |
| oVG109 | TTCTAATACGACTCACTATAGCCGGAATTATTGGGCGTAAAGTTTTAGAGCTAGA |
| oVG110 | TTCTAATACGACTCACTATAGTTTCGCTCGCCGCTACTCAGTTTTAGAGCTAGA |
| oVG111 | TTCTAATACGACTCACTATAGTACACGACGCTCTTCCGATCGTTTTAGAGCTAGA |
| oVG112 | TTCTAATACGACTCACTATAGCGGTGAAATGCGTAGAGATGGTTTTAGAGCTAGA |
| oVG113 | TTCTAATACGACTCACTATAGTGGTCGCCGTATCATTAAAAGTTTTAGAGCTAGA |
| oVG114 | TTCTAATACGACTCACTATAGTGTAGGGAAAGAGTGTCAAGTTTTAGAGCTAGA |
| oVG115 | TTCTAATACGACTCACTATAGCACGAAAGGCGCAACGATCGTTTTAGAGCTAGA |
| oVG116 | TTCTAATACGACTCACTATAGTGAAATGCGTAGAGATGTGGGTTTTAGAGCTAGA |
| oVG117 | TTCTAATACGACTCACTATAGTGCTGATAAGTGTAGATCTGTTTTAGAGCTAGA |
| oVG118 | TTCTAATACGACTCACTATAGAGTGGTGGGTTTCCCCATTGTTTTAGAGCTAGA |
| oVG119 | TTCTAATACGACTCACTATAGCATTTGCTTTCTCTTCCTCGTTTTAGAGCTAGA |
| oVG120 | TTCTAATACGACTCACTATAGCACTGTCTCAACGAGAGACTGTTTTAGAGCTAGA |
| oVG121 | TTCTAATACGACTCACTATAGCAGCCGCGGTAATACGTAGGGTTTTAGAGCTAGA |
| oVG122 | TTCTAATACGACTCACTATAGGGTTGTAGGACACTCTGTAGTTTTAGAGCTAGA |
| oVG123 | TTCTAATACGACTCACTATAGAACCAAGAGGCTTGCCTCTTGTTTTAGAGCTAGA |

**Table S4** SPβ prophage genome comparisons at the gene (CDS) level. No gene is present in only one prophage. The genes not shared by all three are: (i) the *yokH–yokC* island (nuclease/CARD-resistance region), shared by 168 and MB8_B7 and absent from P9_B1 (7 genes); (ii) the thymidylate synthase module, present in MB8_B7 and P9_B1 and absent from 168 (2 genes).

| Prophage origin | Genes | Core (all 3) | Accessory (2 of 3) | Unique (1 of 3) |
|---|---|---|---|---|
| 168 | 180 | 173 | 7 | 0 |
| MB8_B7 | 182 | 173 | 9 | 0 |
| P9_B1 | 174 | 172 | 2 | 0 |

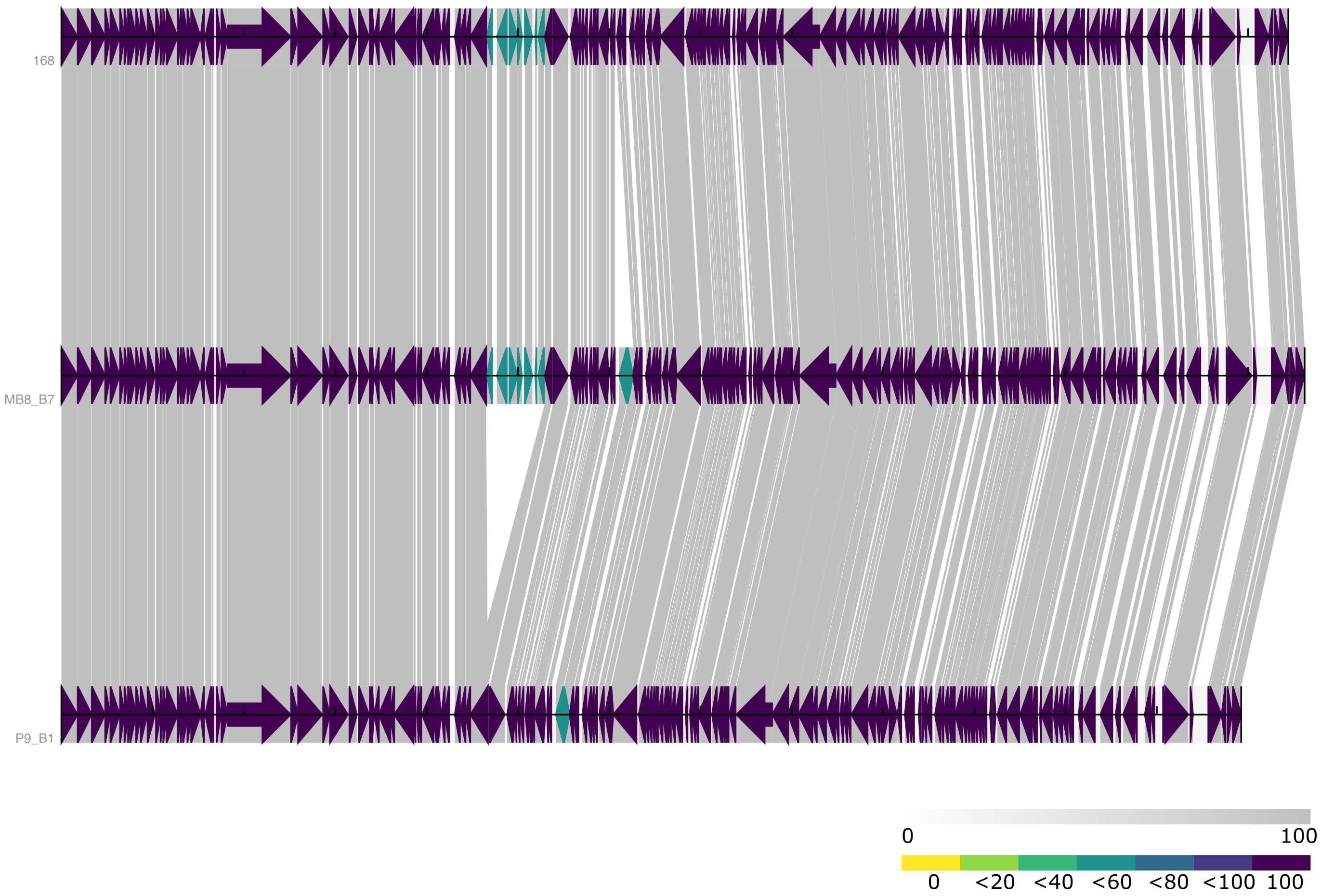


**Figure S1** Representation of the three SPβ prophage genomes. The synteny diagram was obtained with synphage v0.2.7 (Grosboillot and Dragoš, 2024).

**Table S5** Primers used for strain and late competence fluorescence reporter cloning.

| Primer | Sequence (5'->3') |
|---|---|
| amyEF | TCTCCAGTCTTCACATCGGTTTG |
| amyER | GCAAGAGAAAAGTTTTGTCTGATTTATG |
| oAD10 | GAATTCTGCCTGAAAGCTTGTC |
| oAD11 | GCTAGCCATGCTACTGCATCACC |
| oVG126 | CAACGAGCTCCTGCCTTCTCCAAGGAAGG |
| oVG127 | CAACGTCGACGTGACTGTCGGCAATACAC |
| oVG128 | CAACGGATCCTTATTTGTATAGTTCATCCATGCC |
| oVG130 | CAACAAGCTTGCCGGAACTGAATTGGA |

**Table S6** Conditions present in the transcriptomic dataset (for full description of the conditions, see Nicolas et al. (2012)).

| Condition ID | Description |
|---|---|
| aero | growth in synthetic medium until OD578 of 0.5_aerobic conditions |
| B36 | biofilm formation_early biofilm = 36h of growth in MSgg medium without shaking |
| B60 | biofilm formation_strong biofilm = 60h of growth in MSgg medium without shaking |
| BC | 18h of growth on plates_around 1000 colonies per plate |
| BI | 18h of growth on plates_around 100 colonies per plate |
| BMM | growth in BMM medium until OD500 of 0.4_before stress |
| BT | biofilm formation_control for no biofilm = 24h of growth in MSgg medium under shaking |
| C30 | competence_time=30 minutes after induction |
| C90 | competence_time=90 minutes after induction |
| Cold | growth in BMM medium until OD500 of 0.4_60 minutes after shifting from 37 °C to 18 °C |
| dia0 | growth in LB until OD600 of 0.6_before addition of diamide |
| dia5 | growth in LB until OD600 of 0.6_5 minutes after addition of 0.5 mM diamide |
| dia15 | growth in LB until OD600 of 0.6_15 minutes after addition of 0.5 mM diamide |
| Diami | growth in LB until OD540 of 0.3_10 minutes after addition of 0.6 mM diamide |
| Etha | growth in BMM medium until OD500 of 0.4_10 minutes after addition of 4 % ethanol |
| ferm | growth in synthetic medium until OD578 of 0.5_anaerobic coditions in the absence of nitrate |
| Fru | growth in M9 medium until OD600 of 0.5_with D fructose |
| G135 | germination_time=135 minutes after induction |
| G150 | germination_time=150 minutes after induction |
| G180 | germination_time=180 minutes after induction |
| G+S | growth in M9 medium until OD600 of 0.5_with glutamic acid and succinic acid |
| Glu | growth in M9 medium until OD600 of 0.5_with D glucose |
| Glucon | growth in M9 medium until OD600 of 0.5_with gluconate |
| Gly | growth in M9 medium until OD600 of 0.5_with glycerol |
| GM-0 | shift from glucose to malate+glucose_before addition of malate |
| GM+5 | shift from glucose to malate+glucose_5 minutes after addition of malate |
| GM+10 | shift from glucose to malate+glucose_10 minutes after addition of malate |
| GM+15 | shift from glucose to malate+glucose_15 minutes after addition of malate |
| GM+25 | shift from glucose to malate+glucose_25 minutes after addition of malate |
| GM+45 | shift from glucose to malate+glucose_45 minutes after addition of malate |
| GM+60 | shift from glucose to malate+glucose_60 minutes after addition of malate |
| GM+90 | shift from glucose to malate+glucose_90 minutes after addition of malate |
| GM+120 | shift from glucose to malate+glucose_120 minutes after addition of malate |
| GM+150 | shift from glucose to malate+glucose_150 minutes after addition of malate |
| H2O2 | growth in LB until OD540 of 0.3_10 minutes after addition of 0.1 mM H2O2 |
| Heat | growth in BMM medium until OD500 of 0.4_10 minutes after shifting from 37 °C to 48 °C |
| HiOs | growth in SMM medium until OD578 of 1_addition of 1.2 M NaCl |
| HiTm | growth in SMM medium until OD578 of 1_growth temperature of 51 °C |
| HPh | growth in high phosphate medium_exponential phase |
| LBexp | growth in LB medium_exponential phase |
| LBGexp | growth in LB medium with added glucose_exponential phase |

| Condition ID | Description |
|---|---|
| LBGstat | growth in LB medium with added glucose_stationary phase |
| LBGtran | growth in LB medium with added glucose_transition phase |
| LBstat | growth in LB medium_stationary phase |
| LBtran | growth in LB medium_transition phase |
| LoTm | growth in SMM medium until OD578 of 1_growth temperature of 16 °C |
| LPh | growth in low phosphate medium_exponential phase |
| LPhT | growth in low phosphate medium_stationary phase |
| M0t45 | growth in LB until exponential phase_45 minutes after t0 |
| M0t90 | growth in LB until exponential phase_90 minutes after t0 |
| M40t45 | growth in LB until exponential phase_45 minutes after addition of 40 ng/ml mitomycin |
| M40t90 | growth in LB until exponential phase_90 minutes after addition of 40 ng/ml mitomycin |
| M9exp | growth in M9 medium with glucose_exponential phase |
| M9stat | growth in M9 medium with glucose_stationary phase |
| M9tran | growth in M9 medium with glucose_transition phase |
| M+G | growth in M9 medium until OD600 of 0.5_with D glucose and L malic acid |
| Mal | growth in M9 medium until OD600 of 0.5_with L malic acid |
| MG-0 | shift from malate to malate+glucose_before addition of glucose |
| MG+5 | shift from malate to malate+glucose_5 minutes after addition of glucose |
| MG+10 | shift from malate to malate+glucose_10 minutes after addition of glucose |
| MG+15 | shift from malate to malate+glucose_15 minutes after addition of glucose |
| MG+25 | shift from malate to malate+glucose_25 minutes after addition of glucose |
| MG+45 | shift from malate to malate+glucose_45 minutes after addition of glucose |
| MG+60 | shift from malate to malate+glucose_60 minutes after addition of glucose |
| MG+90 | shift from malate to malate+glucose_90 minutes after addition of glucose |
| MG+120 | shift from malate to malate+glucose_120 minutes after addition of glucose |
| MG+150 | shift from malate to malate+glucose_150 minutes after addition of glucose |
| Mt0 | growth in LB until exponential phase_t0 = before addition of mitomycin |
| nit | growth in synthetic medium until OD578 of 0.5_anaerobic coditions in the presence of nitrate |
| Oxctl | growth in LB until OD540 of 0.3_control = 10 minutes growth without oxidative stressor |
| Paraq | growth in LB until OD540 of 0.3_10 minutes after addition of 0.4 mM paraquat |
| Pyr | growth in M9 medium until OD600 of 0.5_with pyruvate |
| S0 | induction of sporulation_time=0 |
| S1 | induction of sporulation_time=1h |
| S2 | induction of sporulation_time=2h |
| S3 | induction of sporulation_time=3h |
| S4 | induction of sporulation_time=4h |
| S5 | induction of sporulation_time=5h |
| S6 | induction of sporulation_time=6h |
| S7 | induction of sporulation_time=7h |
| S8 | induction of sporulation_time=8h |
| Salt | growth in SMM medium until OD500 of 0.4_10 minutes after addition of 0.4 M NaCl |
| SMM | growth in SMM medium until OD500 of 0.4_before stress |

| Condition ID | Description |
|---|---|
| SMMPr | growth in SMM medium until OD578 of 1_growth temperature of 37 °C |
| Sw | 18h of growth on soft plates |
| T-5.40H | batch fermentation in modified M9 medium_340 minutes before glucose consumption |
| T-4.40H | batch fermentation in modified M9 medium_280 minutes before glucose consumption |
| T-3.40H | batch fermentation in modified M9 medium_220 minutes before glucose consumption |
| T-2.40H | batch fermentation in modified M9 medium_160 minutes before glucose consumption |
| T-1.40H | batch fermentation in modified M9 medium_100 minutes before glucose consumption |
| T-1.10H | batch fermentation in modified M9 medium_70 minutes before glucose consumption |
| T-0.40H | batch fermentation in modified M9 medium_40 minutes before glucose consumption |
| T0.0H | batch fermentation in modified M9 medium_time of glucose consumption |
| T0.30H | batch fermentation in modified M9 medium_30 minutes after glucose consumption |
| T1.0H | batch fermentation in modified M9 medium_60 minutes after glucose consumption |
| T1.30H | batch fermentation in modified M9 medium_90 minutes after glucose consumption |
| T2.0H | batch fermentation in modified M9 medium_120 minutes after glucose consumption |
| T2.30H | batch fermentation in modified M9 medium_150 minutes after glucose consumption |
| T3.0H | batch fermentation in modified M9 medium_180 minutes after glucose consumption |
| T3.30H | batch fermentation in modified M9 medium_210 minutes after glucose consumption |
| T4.0H | batch fermentation in modified M9 medium_240 minutes after glucose consumption |
| T5.0H | batch fermentation in modified M9 medium_300 minutes after glucose consumption |

**Table S7** Bacterial genome comparisons at the gene (CDS) level. The 26 unique 168 genes correspond essentially the ICEBs1 element; the 9 unique P9_B1 genes lie in a phage-like region around 2.53 Mb; MB8_B7 has no strictly unique gene.

| Strain | Genes | Core (all 3) | Accessory (2 of 3) | Unique (1 of 3) |
|---|---|---|---|---|
| 168 | 4032 | 3997 | 9 | 26 |
| MB8_B7 | 4003 | 3996 | 7 | 0 |
| P9_B1 | 4010 | 3997 | 4 | 9 |

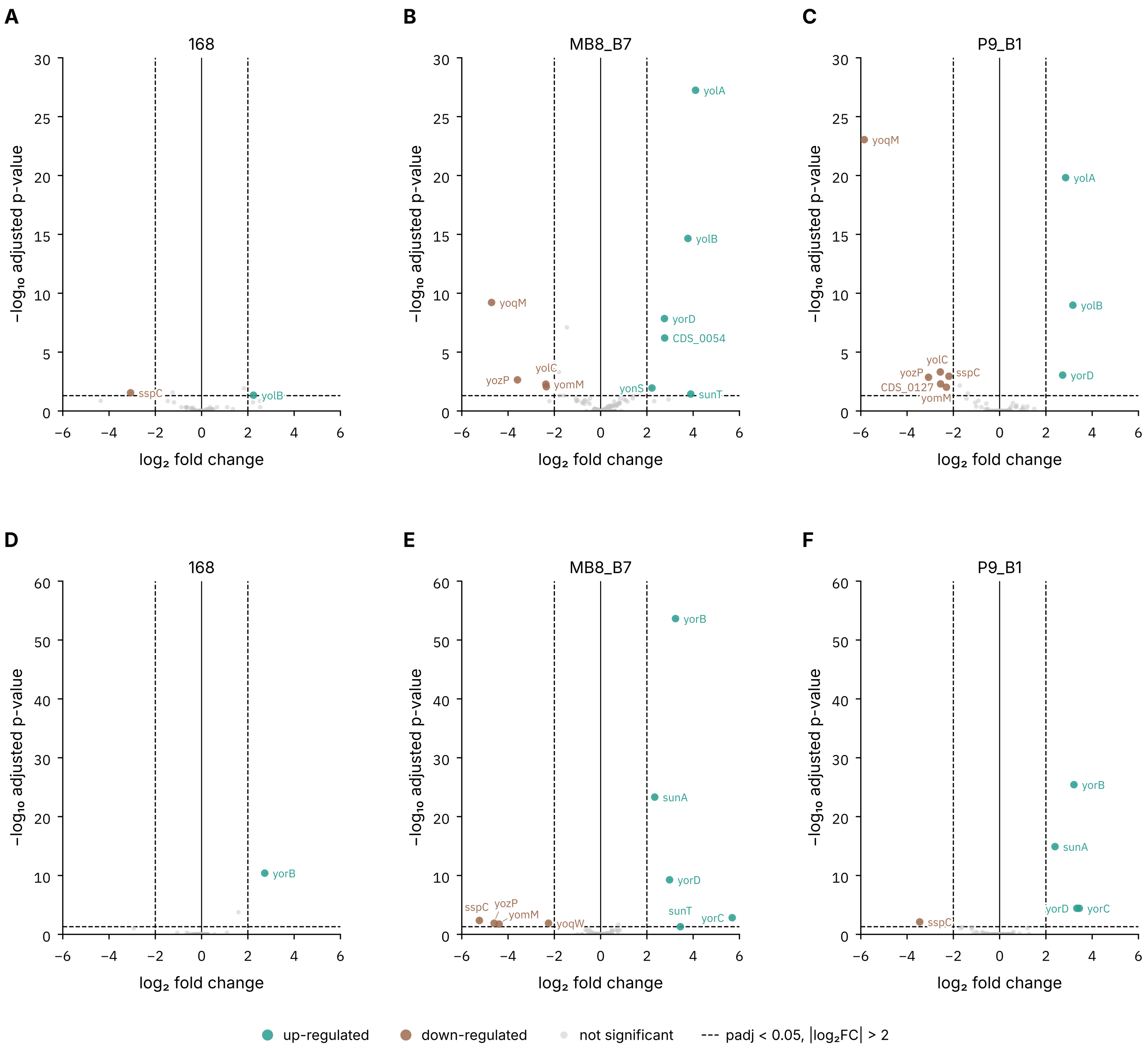


**Figure S2** Prophage gene differential expression between conditions: C90 or M40t90 versus LBstat (lysogens only). The volcano plots using padj < 0.05 and $|\log_2 FC| > 2$ show the prophage core genes affected by the conditions: competence vs. LBstat for 168 (A), MB8_B7 (B) and P9_B1 (C) prophage and mitomycin C vs. LBstat for 168 (D), MB8_B7 (E) and P9_B1 (F) prophage. Upregulated genes are in teal and downregulated genes in brown.

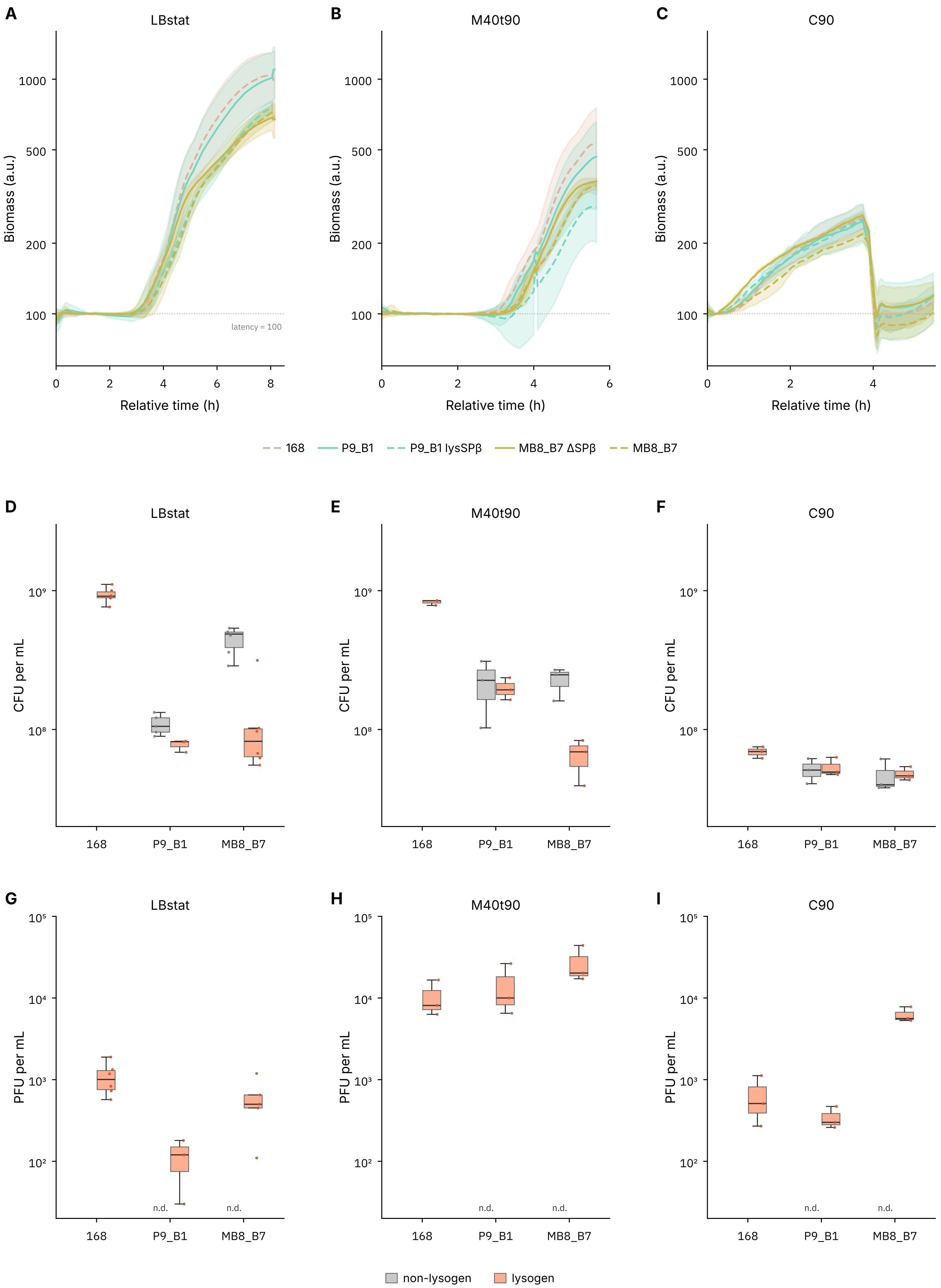


**Figure S3** Growth, bacterial and phage concentrations of RNA-seq samples. Growth curves for LBstat (A), M40t90 (B) and C90 (C) culture conditions. (D, E, F) Respective bacterial concentration in CFU/mL. (G, H, I) Respective phage concentration in PFU/mL. Non-lysogens are shown in grey and lysogens in orange.

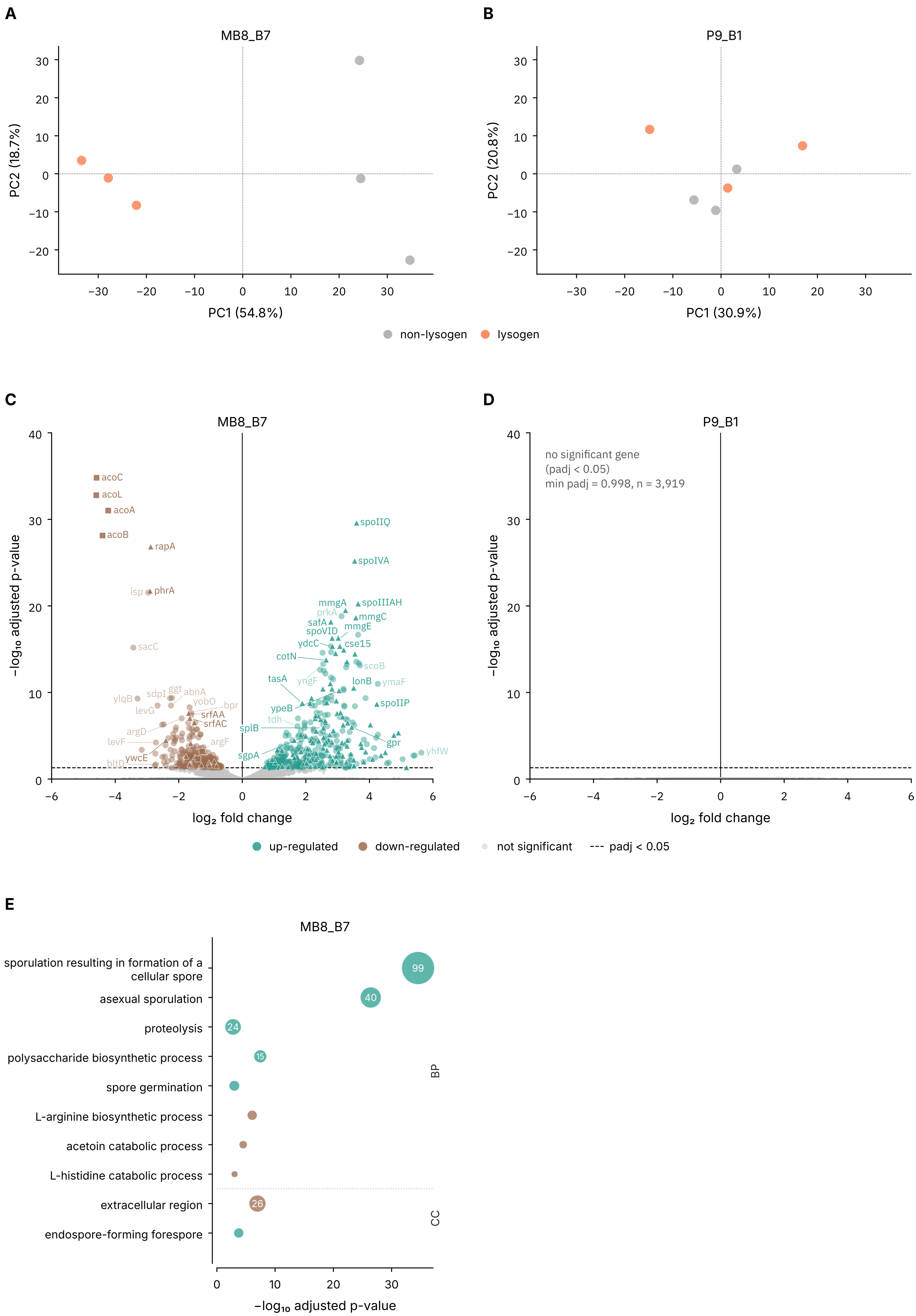


**Figure S4** Differential expression between lysogen vs. non-lysogen (bacterial genes only) during growth in LB to stationary phase. PCA plots for MB8_B7 (A) and P9_B1 (B). Non-lysogens are in grey and lysogens in orange. Volcano plots for MB8_B7 (C) and P9_B1 (D). Significantly upregulated (teal) and downregulated (brown) genes are shown for padj < 0.05. GO enrichment plots for MB8_B7 (E). Sporulation and acetoin catabolism pathway-related genes are highlighted.

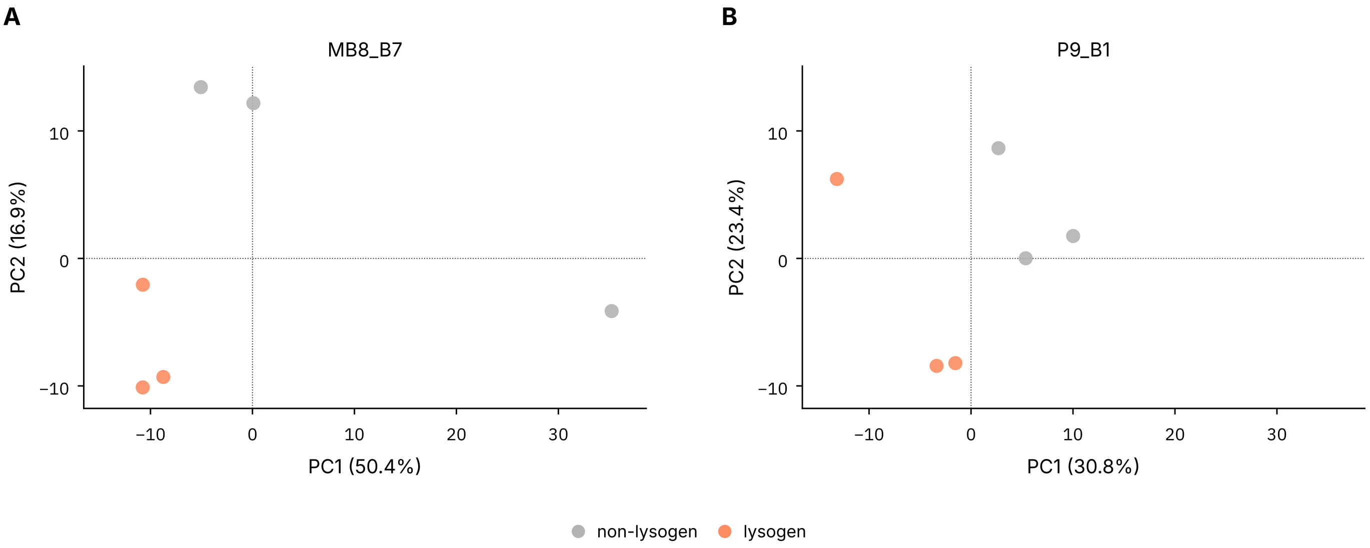


**Figure S5** Differential expression between lysogen vs. non-lysogen (bacterial genes only) during growth in competence medium. PCA plots for MB8_B7 (A) and P9_B1 (B). Non-lysogens are in grey and lysogens in orange.

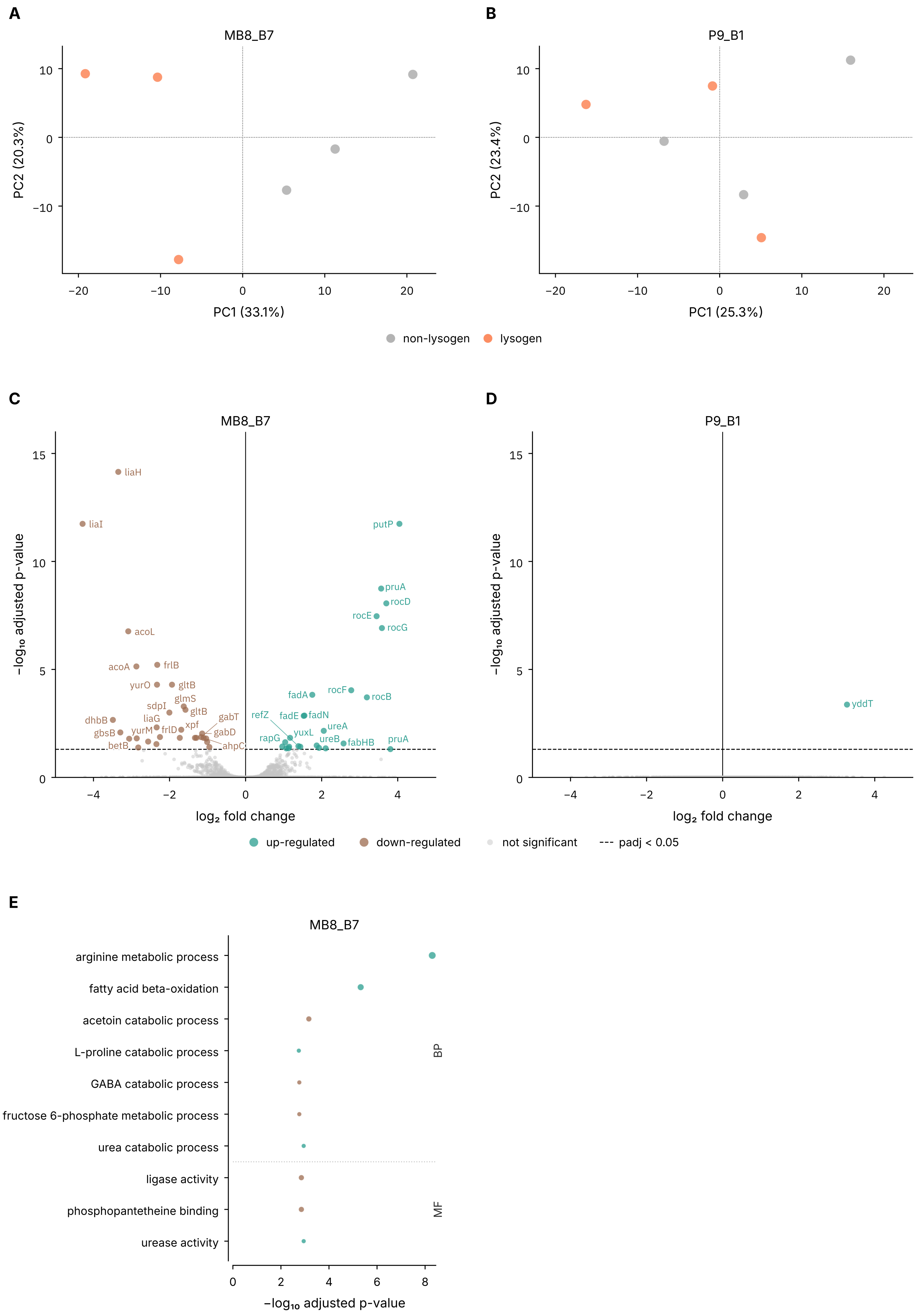


**Figure S6** Differential expression between lysogen vs. non-lysogen (bacterial genes only) during growth in the presence of mitomycin C. PCA plots for MB8_B7 (A) and P9_B1 (B). Non-lysogens are in grey and lysogens in orange. Volcano plots for MB8_B7 (C) and P9_B1 (D). Significantly upregulated (teal) and downregulated (brown) genes are shown for padj < 0.05. GO enrichment plots for MB8_B7 (E).

Grosboillot, V., and Dragoš, A. (2024). Synphage: A pipeline for phage genome synteny graphics focused on gene conservation. *Bioinformatics Advances* 4, vbae126. doi: 10.1093/bioadv/vbae126

Nicolas, P., Mäder, U., Dervyn, E., Rochat, T., Leduc, A., Pigeonneau, N., et al. (2012). Condition-Dependent Transcriptome Reveals High-Level Regulatory Architecture in *Bacillus Subtilis*. *Science* 335, 1103–1106. doi: 10.1126/science.1206848

Wilkins, O. G., and Ule, J. (2021). Ribocutter: Cas9-mediated rRNA depletion from multiplexed Ribo-seq libraries. doi: 10.1101/2021.07.14.451473
